# Spatial architecture of high-grade glioma reveals tumor heterogeneity within distinct domains

**DOI:** 10.1101/2023.03.13.531204

**Authors:** J.J.D. Moffet, O.E. Fatunla, L. Freytag, J.J. Jones, S. Roberts-Thomson, A. Pavenko, D. Scoville, L. Zhang, Y. Liang, A. Morokoff, J.R. Whittle, S. Freytag, S.A. Best

## Abstract

High-grade gliomas are aggressive primary brain cancers with poor response to standard regimens, driven by immense heterogeneity. In isocitrate dehydrogenase (*IDH*) wild-type high-grade glioma (glioblastoma, GBM), increased intra-tumoral heterogeneity is associated with more aggressive disease. Recently, spatial technologies have emerged to dissect this complex heterogeneity within the tumor ecosystem by preserving cellular organization *in situ*. Here, we construct a high-resolution molecular landscape of GBM and *IDH*-mutant high-grade glioma patient samples to investigate the cellular subtypes and spatial communities that compose high-grade glioma using digital spatial profiling and spatial molecular imaging. This uncovered striking diversity of the tumor and immune microenvironment, that is embodied by the heterogeneity of the inferred copy-number alterations in the tumor. Reconstructing the tumor architecture revealed two distinct niches, one composed of tumor cell states that most closely resemble normal glial cells, associated with microglia; and the other niche populated by monocytes and mesenchymal tumor cells. We further reveal that communication between tumor and immune cells is underpinned by tumor-specific ligands, such as TGFβ signaling in astrocyte-like tumor cells. This primary study reveals high levels of intra-tumoral heterogeneity in high-grade gliomas, associated with a diverse immune landscape within spatially localized regions.

## INTRODUCTION

High-grade gliomas, characterized as world health organization (WHO) grades 3 and 4, are aggressive forms of brain cancer with poor survival outcomes and treatment regimens that have not changed in decades. The evolution of WHO grading^1^ has increasingly incorporated molecular analysis that stratifies glioma based on isocitrate dehydrogenase (*IDH*) status into groups with distinct biology and clinical outcomes with mutations in *IDH1/2* predicting improved survival (5 – 15 years, depending on grade)^2–5^. *IDH* mutated gliomas are further stratified into oligodendroglioma which is defined by the presence of 1p19q co-deletion and astrocytoma (1p19q intact). *IDH* wild type status characterizes grade 4 glioblastoma (GBM) with a median prognosis with treatment of just 13-18 months^6^. Multiple genetic alterations are additionally used to classify gliomas, including *ATRX* loss, deletions or inactivating mutations in *TP53, PTEN*, *NF1*, and *CDKN2A/B*, amplification of *EGFR*, *PDGFRA*, and *CDK4/6,* and gain of chromosome 7 and loss of 10 ^1,7^. While molecular profiling provides important prognostic value, novel therapies that target the molecular drivers of high-grade glioma have yet to show clinical benefit^8,9^. Thus, patients still receive radiation and alkylating chemotherapy^4,5,10,11^ using regimens unchallenged for the past 25 years, highlighting a need for improved stratification and treatment options.

In addition to genetic alterations, intrinsic cellular plasticity adds to the intra- and inter-patient phenotypic variability observed in glioma^12^. Advances in single cell sequencing revealed that tumor cells exhibit transcriptional signatures mirroring early development of the healthy human brain; including astrocytic, oligodendrocytic, neural, mesenchymal and highly proliferative progenitor cell states^13–17^. *IDH*-mutant high-grade glioma displays a bi-lineage hierarchy of astrocytic and oligodendrocytic cell states, with a rare subpopulation of progenitor cells^16,17^, defining astrocytoma and oligodendroglioma tumors. In comparison to *IDH*-mutant high-grade glioma, GBM tumors are characterized by greater intra-tumoral heterogeneity, the extent correlating with survival^18^, resulting in a mixture of each cell state within the tumor^13,14,18^. Upon recurrence, there is a shift towards the mesenchymal lineage likely induced by immune cells and interferon signaling^19–21^. The immense plasticity of glioma tumor cells allows them to reprogram and evade treatment, contributing to the poor survival of patients^22^.

Overcoming tumor heterogeneity is a major challenge for cancer treatment, with the past decade focusing on novel immunotherapies that drive an antigen-mediated anti-tumor response, determined by the composition and extent of immune infiltration. While these strategies have been successful in many solid cancers, particularly those with high tumor mutational burden, to date efficacy in glioma has been limited. A number of factors likely contribute to the lack of benefit in glioma. Firstly, the immune microenvironment of the brain is classically associated with poor infiltration, with an immunosuppressive environment and largely composed of resident microglia cells^23^. Indeed, both *IDH*-mutant high grade and GBM tumors are dominated by myeloid cells, though GBM tumors exhibit increased bone marrow derived macrophages (BDM) and lymphocytes compared to *IDH*-mutant tumors^24^. However, there have been reported association of tumor cell states and specific immune cell infiltrate, such as mesenchymal cells with myeloid^14,20^ and T cell infiltration^25^, which were associated with a more unfavorable prognosis^20^. Tumor cells together with the immune microenvironment create a complex milieu that ultimately promotes adaptability and disease progression, highlighting the need for a detailed understanding of the interactions between immune cells and different tumor cell states.

To capture a deeper understanding of the tumor microenvironment of high-grade gliomas, we perform a combination of digital spatial profiling (DSP)^26^ and spatial molecular imaging (SMI)^27^. In concert, these technologies, which preserve the tissue architecture, uncover the overall identity of the spatial environment^28^ with complementary strengths. Sequencing based technologies capture regions of tissue to provide next-generation sequencing (NGS) output of the whole transcriptome, and while not providing single-cell resolution, these technologies are not limited by gene number. In contrast, high-plex imaging technologies using a probe-based readout offer subcellular resolution, but are currently limited to a restricted number of gene targets. Integrating these technologies, we confirm immense spatial heterogeneity in GBM samples compared to *IDH1*-mutant tumors, characterized by the presence of the mesenchymal tumor cell lineage. Through the isolation and analysis of cycling Ki67^+^ tumor cells, we find that progenitor cells emerge within this subset, highlighting the plasticity of glioma cell states. Analysis of the immune compartment revealed discrete neighborhoods with patterns of tumor and immune cell interaction that validated across GBM cohorts and independent spatial sequencing technologies. Finally, we create a map of tumor cell state and immune cell interaction in the GBM microenvironment. Our findings highlight the immense heterogeneity of tumor cell states in GBM and their relationship with the immune microenvironment.

## RESULTS

### Spatial transcriptomics analysis of high-grade glioma

To interrogate the spatial intra-tumoral heterogeneity of high-grade glioma, we selected three *IDH*-wild type WHO grade 4 GBM and three *IDH1-*mutant grade 3 astrocytoma (A) samples for spatial whole transcriptome analysis. Pathology review of the six cases found these were representative and showed diagnostic morphological features, with astrocytoma samples displaying homogenous pleomorphic cells and glioblastomas demonstrating a higher degree of anaplasia (**Table 1, Fig. 1a**). Regions of interest (ROI) were selected based on histological features, distribution of proliferating tumor cells and areas of central tumor (CT) bordering on infiltrating tumor (IT) and leading edge (LE) regions across the six samples (**Supplementary Fig. 1a**). By immunostaining for GFAP, CD45 and Ki67 protein expression, 34 areas of illumination (AOI) representing tumor cells (T, GFAP^+^), 10 AOIs representing proliferating tumor cells (K, GFAP^+^Ki67^+^) and 20 AOIs representing immune cells (I, CD45^+^) were isolated within each ROI using Digital Spatial Profiling (DSP) on the NanoString GeoMx^®^ platform (**Table 2, Supplementary Fig. 1b, Supplementary Tables 1 and 2**).

**Fig. 1.**
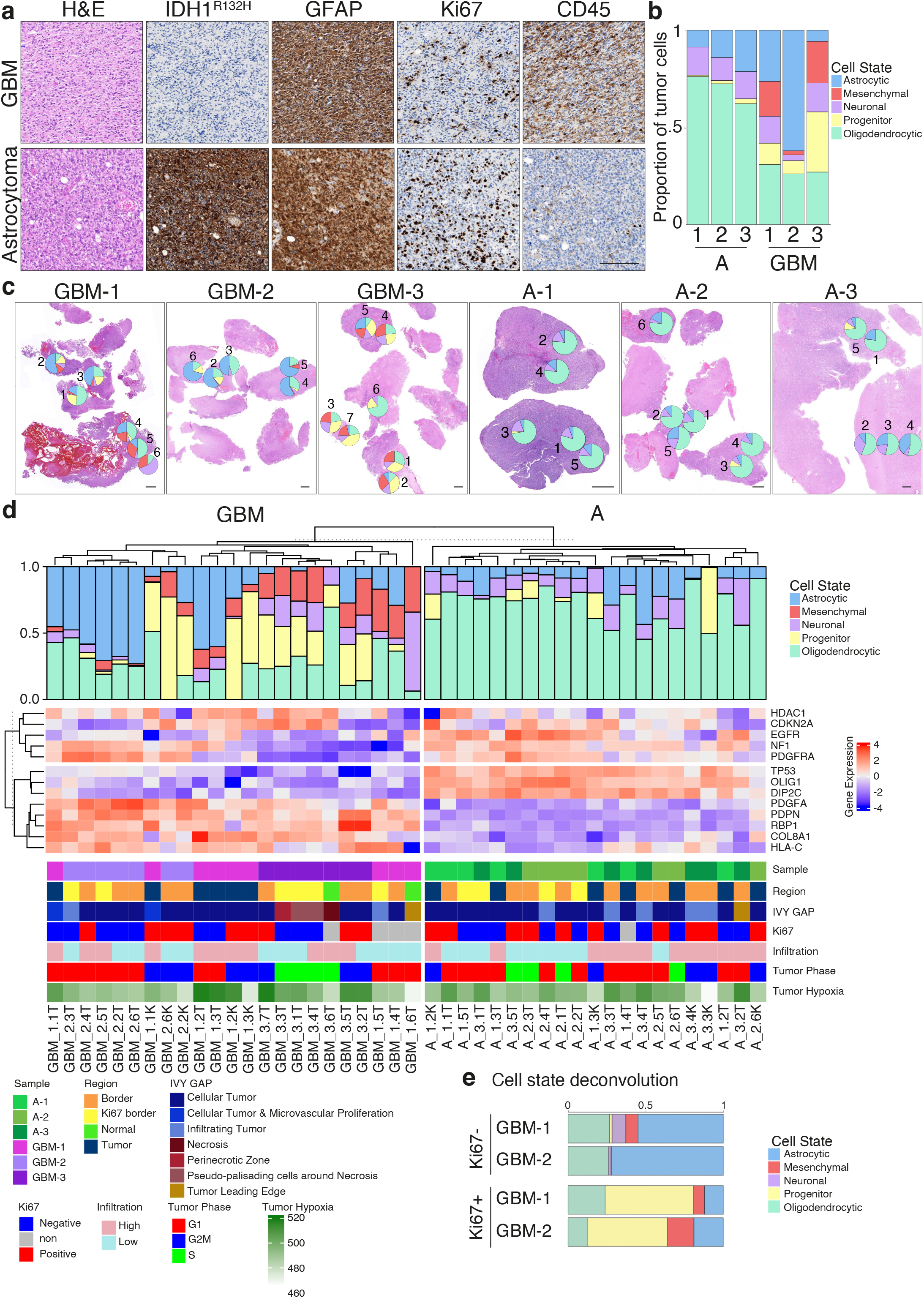
Detecting regions of interest in high-grade glioma. **a.** Immunostaining of GFAP, IDH1-R132H, Ki67, CD45 and H&E stain of representative regions in GBM and astrocytoma (A) samples. Scale, 200 µm. **b.** Deconvolution of tumor cell states per patient sample (A n=3, GBM n=3). **c.** H&E of GBM and astrocytoma samples with tumor cell state proportions plotted over each region of interest (ROI). Scale, 1 mm. **d.** Complex heatmap of each tumor (T) and Ki67^+^ tumor (K) AOI deconvolution and key characteristics. Rows and columns are clustered by expression of selected genes, following splitting into GBM (left) and astrocytoma (right) samples, and splitting into clinically relevant (above) and differentially expressed (below) genes. Above the heatmap, columns are annotated with the tumor deconvolution of each AOI; below, columns are annotated for each AOI patient sample of origin, location of ROI in relation to tumor immunostaining, pathology classification based on Ivy GAP, Ki67 status based on immunohistochemistry, level of immune infiltration based on CD45 immunohistochemistry, cell cycling phase based on gene expression, and tumor hypoxia based on gene expression. **e.** Deconvolution of GBM-1 (n=3) and GBM-2 (n=2) Ki67^-^ vs Ki67^+^ AOI in CT regions. Proportions averaged across AOI.

**TABLE 1.** Characteristics of patient samples.

| Patient information |  |  |  | Clinical information |  |  |  |  |  |  |  |  |
| --- | --- | --- | --- | --- | --- | --- | --- | --- | --- | --- | --- | --- |
| Sample | Age | Sex | Tumor location | IDH1 | ATRX | TP53 | CDKN2A | Ki67 | CD45 | Cytology | WHO grade | Comments |
| A-1 | 35 | F | Left frontal | Mutated | Intact | Intact | Intact | 8-13% | – | A* | 3 | Nil |
| A-2 | 27 | F | Right parietal | Mutated | Mutated | Mutated | Intact | 18-23% | + | A | 3 | Pre-treated: radiation |
| A-3 | 45 | M | Right frontal | Mutated | Mutated | Mutated | Mutated | 10-15% | + | A | 3 | Pre-treated: radiation, temozolomide |
| GBM-1 | 67 | M | Right temporo-occipital | Intact | Intact | Mosaic | Intact | 10-15% | – | GBM | 4 | MGMT methylated |
| GBM-2 | 49 | M | Left frontal | Intact | Intact | Intact | Mutated | 10-15% | ++ | GBM | 4 | MGMT unmethylated |
| GBM-3 | 47 | M | Left temporal | Intact | Intact | Mosaic | Mutated | 20-30% | + | GBM | 4 | MGMT unmethylated |

GBM, glioblastoma; A, astrocytoma; F, female; M, male.
\* Sample A-1 was re-diagnosed to an oligodendroglioma following the detection of 1p/19q codeletion by next generation sequencing panel

**TABLE 2.** GeoMx^®^ DSP spatial transcriptomics key features.

| Features |  | A | GBM |
| --- | --- | --- | --- |
| Sample | Patients | 3 | 3 |
|  | ROI | 16 | 18 |
|  | AOI | 32 | 39 |
|  | Excluded | 6 | 1 |
| AOI | Tumor | 16 | 18 |
|  | Ki67+ | 5 | 5 |
|  | CD45 | 5 | 15 |

To analyze the tumor cell compartment, we first combined the bulk transcriptomic data from the tumor AOIs (T combined with K for the same ROI). Principal component analysis (**Supplementary Fig. 1c, d**) and deconvolution into the core tumor cell states^13^ (**Supplementary Fig. 2a**) reveal large differences between astrocytoma and GBM; most notably the increase of mesenchymal (p<0.001) and progenitor (p<0.001) tumor cell states in GBM compared to astrocytoma (**Fig. 1b**). Further investigation of the tumor states within each ROI revealed spatial heterogeneity with significant increases in intra-tumoral heterogeneity in the GBM samples compared to astrocytoma in the mesenchymal (variance test, p<0.001) and progenitor (variance test, p<0.001) tumor cell populations (**Fig. 1c**). As expected, proliferating (K) AOIs were significantly enriched with progenitor tumor cell states compared to AOIs of non-proliferating tumor (T) AOIs (26.2% and 4.5% of estimated nuclei respectively, p<0.001) (**Fig. 1e**). This finding is consistent with the association of proliferation and the glioma stem cell (GSC) with the progenitor subtype in previous studies^13^ and a significant portion of the progenitor cell signature contained key cycling genes (including *TOP2A, MKI67, AURKB*) (**Supplementary Fig. 2b**). Hierarchical clustering of all tumor AOIs did not seem to be driven by patient sample, histological classification, cycling state, immune infiltration of the region nor hypoxic status in this small cohort (**Fig. 1d**), suggesting that the heterogeneity of these glioma samples may not be associated by macroscopic events.

### Spatially localized tumor cells display distinct chromosomal alterations

Utilizing the whole transcriptome sequencing data, we applied CNVkit^29^ to investigate chromosomal regions containing under- or over-represented housekeeping genes, to predict chromosomal alterations. For the GBM and *IDH1*-mutant glioma samples, 7.5 % and 70.5 % of all copy number alterations (CNA) beyond two standard deviations from the mean, were heterogeneous between patients, respectively; however, the majority of CNAs were driven by both inter- (between) and intra-tumoral (within) heterogeneity (**Supplementary Fig. 3a, b**). In the *IDH1*-mutant Astrocytoma samples, intra-tumoral heterogeneity was identified in chromosomal regions involved in tumor promotion and immune evasion (*NOTCH1, TP53I3, HLA*) (**Fig. 2a**). Inter-tumoral heterogeneity was identified significantly in chromosome arm 1 p, driven by A-1, and associated with loss of chromosome 19 q, indicative of an oligodendroglioma^30^ (**Fig. 2b**). Further examination of the intra- and inter-tumoral gene expression heterogeneity across the astrocytoma samples clearly indicated that the oligodendrocytic cell state was a key driver of heterogeneity between astrocytoma samples, independent of cell cycling state (**Fig. 2c**). Follow-up with the treating hospital confirmed that patient A-1 had received an updated diagnosis after next generation sequencing, thus independently validating our CNVkit CNA analysis on DSP technology. Partitioning the heterogeneity of only A-2 and A-3 demonstrated disposition towards intra-tumoral over inter-tumoral heterogeneity driven by the proliferating progenitor tumor state (**Supplementary Fig. 4a**). Differential gene expression analysis between the oligodendroglioma and astrocytoma samples revealed an enrichment of genes involved in neural signaling pathways in oligodendroglioma (including *IGFBP3* ^31,32^, *KIF3C* ^33,34^ and *SCG2* ^35,36^) (**Supplementary Fig. 4b, c**), however future analysis with a larger sample size will be required to investigate these aspects further.

**Fig. 2.**
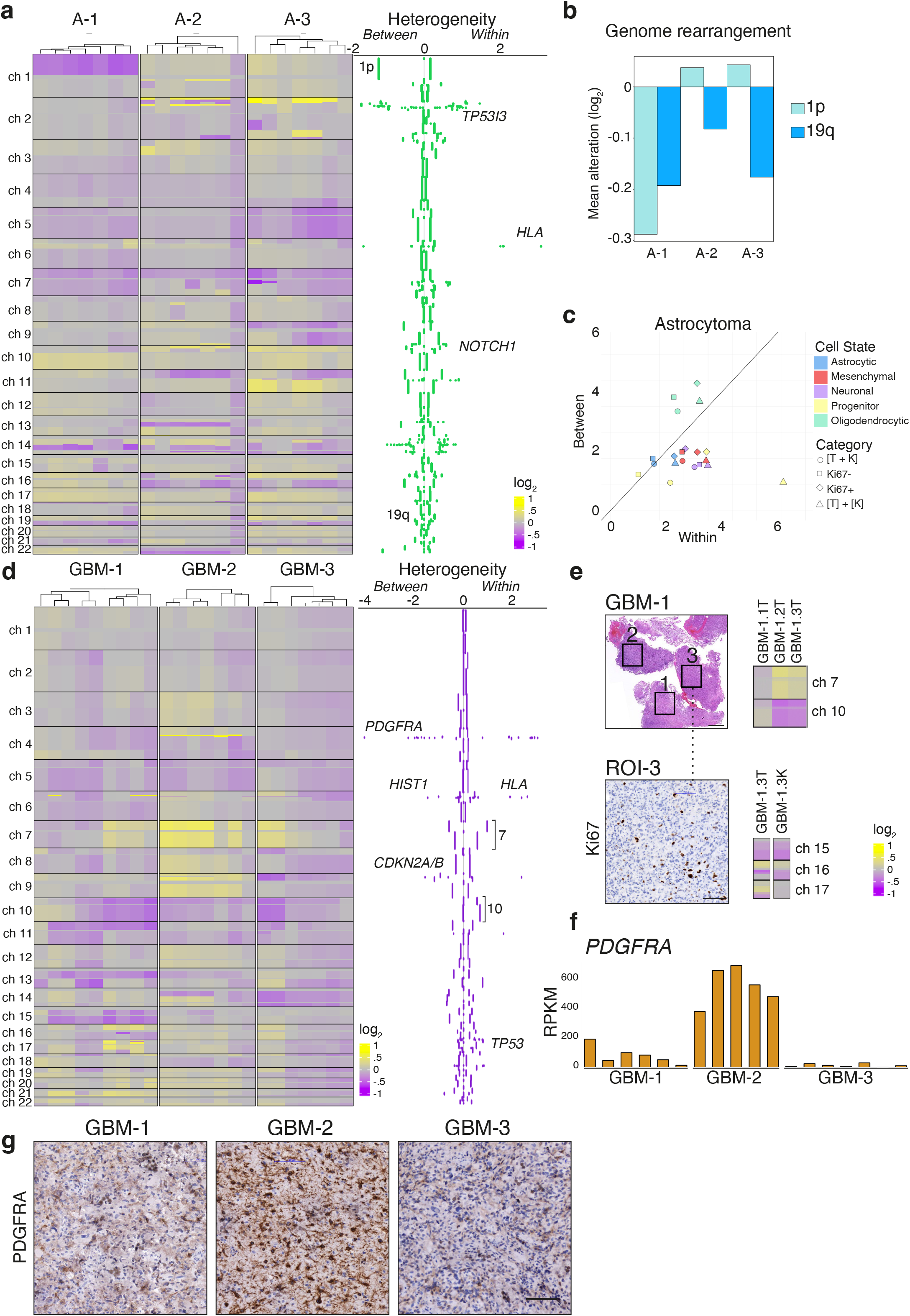
Spatial heterogeneity detected by copy number alteration. **a.** Copy number alteration (CNA) analysis per chromosome of astrocytoma samples, with heterogeneity overview. Left, inferred CNA segmentation, displaying windows of 10,000 base pairs. Right, between sum of squares and within sum of squares values for each window, log_2_-transformed. Sum of squares was calculated using only astrocytoma samples. **b.** Chromosome arms 1p and 19q comparison across astrocytoma samples. Average CNA value across chromosome arm, compared to average CNA value across the whole genome. **c.** Shannon Entropy analysis of cell state in astrocytoma tumor samples. Colored by cell state, and repeated for both combined ([T + K]) and separated ([T] + [K]) datasets, as well as Ki67^-^ and Ki67^+^ samples only. **d.** CNA analysis per chromosome of GBM samples, with heterogeneity overview. Left, inferred CNA segmentation, displaying windows of 10,000 base pairs. Right, between sum of squares and within sum of squares values for each window, log_2_-transformed. Sum of squares was calculated using only GBM samples. **e.** GBM-1 ROI 1-3 location (left) and CNA plots depicting chromosome 7 and chromosome 10 (right). ROI 3 (below) Ki67 immunostaining and GFAP^+^Ki67^+^ (K) and GFAP^+^Ki67^-^ (T) CNA plots depicting chromosomes 15-17. **f.** *PDGFRA* gene expression in GBM samples (n=3). **g.** Immunohistochemistry of PDGFR alpha expression in GBM samples. Scale, 100 µm.

Next, investigating the GBM patient samples, the highest inter-tumoral heterogeneity in terms of CNAs was identified in GBM-2, whilst the highest intra-tumoral heterogeneity was found between the ROIs in GBM-1 (**Supplementary Fig. 3a, b**). Across all GBM samples, the majority of ROIs displayed the prominent GBM genetic alteration, gain of chromosome 7 and loss of chromosome 10 (**Fig. 2d**) further confirming the validity of the CNVkit approach. Both spatial distribution and proliferation appear to play a role in the CNA heterogeneity within samples. For example, in GBM-1 three ROIs in close proximity display heterogeneity in gain of chromosome 7 and loss of chromosome 10 (**Fig. 2e**), reflected in substantial cell state heterogeneity between the regions (**Fig. 1c**). Furthermore, within region 3 of GBM-1, proliferative and non-proliferative AOI also display substantial CNA heterogeneity on chromosomes 15-17 (**Fig. 2e**). To further examine tumor heterogeneity, we examined the impact of the tumor state and proliferative status on the GBM samples using a Shannon entropy approach (**Supplementary Fig. 5a**). This analysis finds that the tumor states of non-proliferating tumor samples drive inter-tumoral heterogeneity, while the proliferating (Ki67^+^) samples mostly contribute towards intra-tumoral heterogeneity. Intra-tumoral entropy was highest when looking at the progenitor signature among separated T and K AOIs (3.2 standard deviations from the overall mean intra-tumoral entropy score). This evidence suggests that the proliferative tumor cells drive a significant phenotypic divide within samples. Examination of the gene expression heterogeneity (**Supplementary Fig. 5b, c**) finds genes associated with the astrocytic (*APOE, AQP4, GFAP*) and oligodendrocytic (*PDGFRA*) lineages driving heterogeneity between tumors and mesenchymal-associated genes (*CHI3L1, VEGFA*) driving intra-tumoral heterogeneity. Notably, the heterogeneity seen in *PDGFRA* expression across patients is mirrored by a focal gain in chromosome 4 in GBM-2, with resulting inter-tumoral differences in RNA and protein expression (**Fig. 2f, g**). To validate our findings in an independent dataset, we interrogated a recently deposited GBM Visium^®^ spatial transcriptomics cohort^15^. Visium^®^ technology analyzes the whole transcriptome from 55 µm spots^28^. The lipid transporter Apolipoprotein E (*APOE*), identified as highly associated with inter-tumoral heterogeneity in our data (**Supplementary Fig. 5b**), similarly displayed homogenous expression within individual GBM samples, yet showed variable expression between samples in the Visium^®^ dataset (**Supplementary Fig. 5d**). Furthermore *VEGFA*, a variable gene within patient samples (**Supplementary Fig. 5b**), was similarly found to occur in patches in the independent GBM samples, highly correlating with the mesenchymal signature (**Supplementary Fig. 5e**). Consistent with recent single-cell studies^19,37^, our findings point to individual CNAs leading to overall heterogeneity of patient’s transcriptional profiles, while proliferation status determines intra-tumoral differences.

### Glioblastoma architecture reveals distinct cell state niches

To further dissect the cellular identities and interactions of individual cells within the regions of interest, we performed Spatial Molecular Imaging (SMI) of 1,000 RNA probes at subcellular resolution^27^ on sample GBM-1 using the NanoString CosMx^®^ platform. Fields of view (FOV) for SMI analysis were selected based on the location of ROI in the initial DSP analysis, with seven FOV selected across the tumor (**Fig. 3a**). From the 1,000 RNA probes, we calculated sufficient gene distribution to assign tumor cell states and immune cell types of interest for the study (**Supplementary Fig. 6a**). A total of 11,801 cells were assigned, containing 1,978,131 total transcripts with a mean of 168 molecules expressed per cell. These cells were clustered identifying 35 clusters (**Supplementary Fig. 6b**), which were first annotated by tumor, vasculature, or immune cells (**Fig. 3b**). Unsupervised prediction of cell types identified 21 unique cell types of the tumor microenvironment (**Supplementary Fig. 6c**), highlighting major glial and immune cell types predicted in the tumor. To better characterize the tumor cell compartment relative to the initial DSP study, the tumor cells were re-clustered and reannotated using unsupervised prediction according to the five cell lineages^13^ (**Fig. 3c**). Assignment of these clusters to groupings based on our available data from SMI analysis resulted in five updated clusters: mesenchymal; astrocytic; a progenitor cluster comprising of neural progenitor cells (NPC), oligodendrocyte progenitor cells (OPC) and radial glia (RG); oligodendrocytic (mature, normal-like); and one mixed population of tumor cells (**Fig. 3d**). As expected, the progenitor cluster was the most transcriptionally active, three-fold higher than the astrocytic cell lineage (**Fig. 3e, f**). To further investigate the mixed population, we identified the top differentially expressed genes compared to every other cluster and found that cells in the mixed cluster are characterized by a strong down-regulation of *VIM*, *CLU* and *SERPINA3* (**Supplementary Fig. 7a, b**). Consistent with this differential expression, protein expression of Vimentin was moderate throughout GBM-1, further suggesting a population of tumor cells absent in the marker (**Supplementary Fig. 7c**). Investigation of scRNAseq data from independent datasets harmonized from 26 glioblastoma studies^14^ confirmed that regions of low *VIM* expression co-localized with cells matching the signature of the mixed cluster (**Supplementary Fig. 7d**). To determine the identity of the population, upregulated genes were examined relative to the other cell states, finding that gene programs associated with chemokines and signaling to be enriched in the population, suggesting these cells may create an immune-responsive niche in the tumor (**Supplementary Fig. 7e, f**). Downregulated pathways highlighted differentiation from the neural progenitor lineage (**Supplementary Fig. 7g**), together suggesting that this cluster may represent cells differentiating towards a more mesenchymal-like cell fate.

**Fig. 3.**
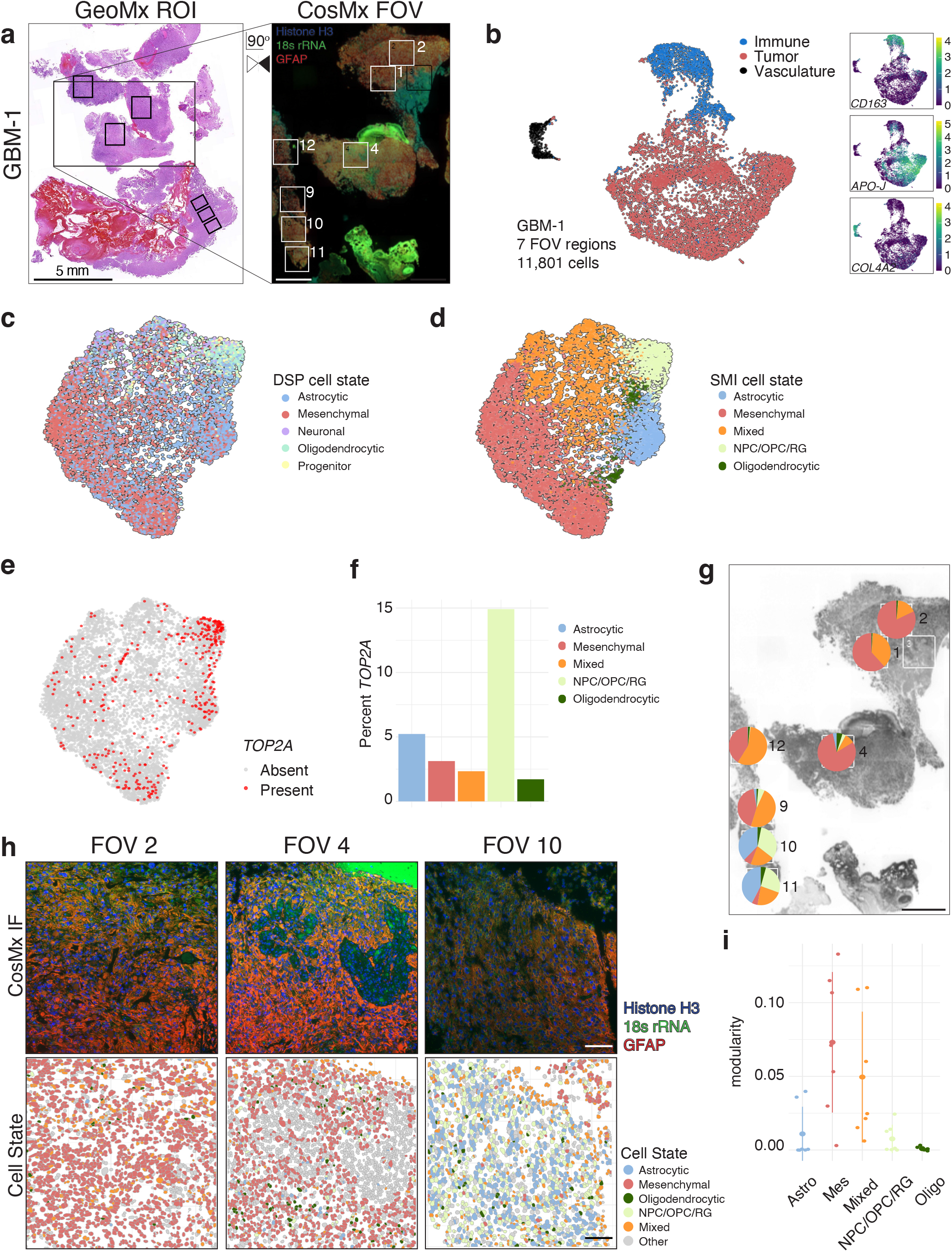
Spatial heterogeneity at the single cell level in glioblastoma. **a.** Schematic of GBM-1 with ROI from GeoMx^®^ DSP analysis (left) and fields of view (FOV) from CosMx^®^ SMI analysis. Scale, 1 mm. **b.** UMAP plot depicting 11,801 cells separated in tumor, immune, and vasculature cell types. Insets show UMAP colored by log expression of marker genes, *APO-J*, *CD163,* and *COL4A2*. **c.** UMAP plot depicting 8,974 tumor cells colored by predicted DSP tumor states based on Couturier *et al.* **d.** UMAP plot of tumor cells annotated by identified tumor states. **e.** UMAP depicting presence of *TOP2A* expression. **f.** Percentage of cells expressing *TOP2A* in each tumor state across all FOVs. **g.** Proportions of each tumor cell state as estimated from the SMI in GBM-1 FOV regions. Scale, 1 mm. **h.** Overview of representative FOV 2, 4, 10 SMI immunofluorescence protein expression (Histone H3, 18s rRNA, GFAP) and Voronoi plots showing cell segmentation and colored by tumor state assignment. Scale, 100 µm. **i.** Modularity score for each tumor cell state, measuring the number of connections between cells of the same state compared to those to other states. Each point indicates one FOV in GBM-1.

To examine the spatial distribution of tumor states, we calculated the overall composition in each FOV. This identified regions enriched with mesenchymal cells, in close proximity to those enriched in astrocytic and progenitor cells (**Fig. 3g**), similar to the degree of spatial heterogeneity observed in the DSP study (**Fig. 1c**). Examining the spatial architecture at a single cell level revealed well-defined patterns of mesenchymal patches and diffuse regions of astrocytic and progenitor cell types (**Fig. 3h, Supplementary Fig. 8a, b**). To quantify the structures of the different tumor states, we performed a modularity analysis, which measures the purity of localized patches of a certain type. As observed visually, the mesenchymal and mixed tumor populations cluster by themselves more than cells of astrocytic (versus mesenchymal p-value = 0.01, versus mixed p-value = 0.06), oligodendrocytic and progenitor tumor states (versus mesenchymal p-value = 0.01, versus mixed p-value = 0.04) (**Fig. 3i**), which co-occur in tumor regions. Therefore, two tumor niches emerge from the spatial data: a brain-native niche derived of intermixed tumor cell states with direct transcriptional and phenotypic relationship with their glial cell-of-origin, comprised of astrocytic, oligodendrocytic and neural progenitor tumor cells; and a second niche, containing tightly packed tumor cells of mesenchymal and mixed phenotype, that have markers suggesting enhanced interaction with immune cells.

### Immune infiltration is associated with the tumor niche

The brain has a distinct immune composition within the tumor microenvironment, largely consisting of Tumor Associated Macrophages (TAMs) - microglia (MG) and bone marrow derived macrophages (BDM), with poor infiltration of lymphocytes^23^. Investigation of the immune compartment across the GBM samples from the DSP data revealed variation in CD45^+^ infiltration, and an enrichment of CD68^+^ TAMs in GBM-1 and GBM-2 (**Fig. 4a**). Consistent with the immunostaining, immune deconvolution of the CD45^+^ immune AOI (**Supplementary Fig. 9a**) revealed an enrichment in microglia and macrophages in GBM-1 and 2, while in GBM-3 a substantial increase in lymphocytes and neutrophils was identified (**Fig. 4b**). While the immune AOIs were contaminated with neoplastic cells (**Supplementary Fig. 1d**), immune cell proportions were unchanged independent of the addition of neoplastic cell types to the deconvolution reference (**Supplementary Fig. 9b, c**). Further deconvolution by region of interest revealed a correlation between the astrocytic lineage and TAMs (correlation: 0.72) and the mesenchymal population with lymphocytes and neutrophil inflammation (correlation: 0.58) (**Fig. 4c**). Enrichment in BDMs associated with the expression of macrophage gene *AHR*, while microglia associated with the expression of *P2RY12* and *TMEM119* (**Supplementary Fig. 9d**). To dissect the immune subtypes and location at a single-cell level, the immune cluster in the SMI analysis of GBM-1 (**Fig. 3b**) was extracted and re-clustered (**Fig. 4d, Supplementary Fig. 10a**). Gene expression analysis revealed a cluster with dominant expression of *AHR,* denoting the identification of BDMs (**Fig. 4e**), opposed to microglia. Taken together, the SMI immune analysis revealed a predominance of macrophages and microglia, with few monocytes and T cells (**Fig. 4d**). The proportions of immune cells in GBM-1 from the deconvolution DSP analysis and single cell SMI analysis revealed similar predictions from each technology, validating both our analysis approaches (**Fig. 4f**). Next, we investigated the spatial distribution of the key immune cell types present in GBM-1, with FOV 2, 4 and 10 having the highest proportion of monocytes, BDMs and microglia, respectively (**Fig. 4g**). When present, monocytes generally distribute in dense clusters of cells (average distance to each other 185 ± 57 µm), while microglia (average distance to each other 356 ± 34 µm) and BDMs (average distance to each other 347 ± 50 µm), were more broadly dispersed throughout the tissue. Thus, the cooccurrence of immune cells with other immune and tumor cell states may be a key factor in the localization of these myeloid cells.

**Fig. 4.**
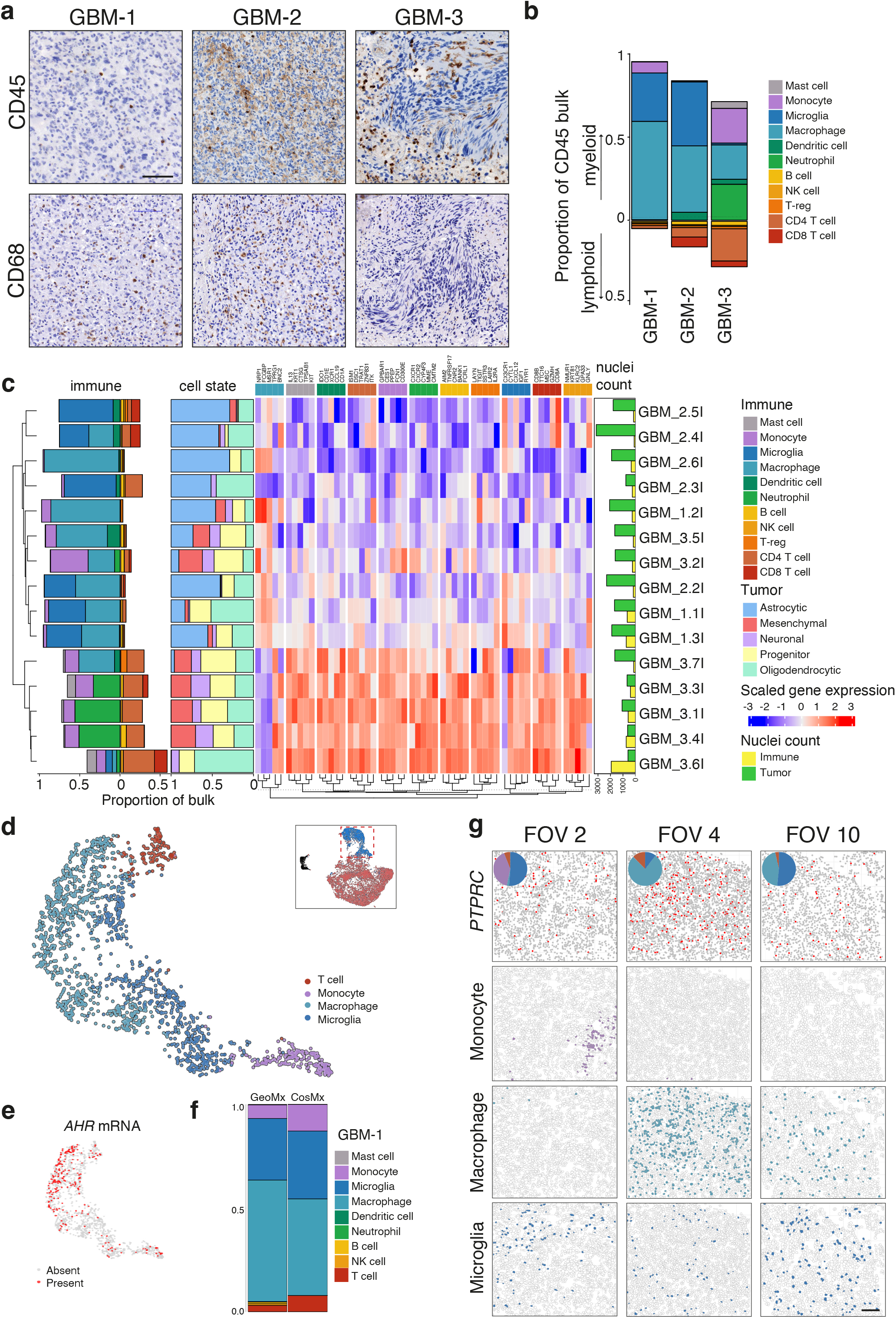
Immune infiltration in glioblastoma. **a.** IHC staining of CD45 and CD68 protein expression of GBM-1 ROI 1, GBM-2 ROI 6 and GBM-3 ROI 7. Scale, 100 µm. **b.** Average proportion of myeloid and lymphoid immune infiltrate in GBM ROIs for each sample, as determined from deconvolution. **c.** Complex heatmap of immune and cell state deconvolution, including only GBM samples with matched immune and tumor AOIs in combined dataset (n = 15). The top 5 genes with highest signature scores for each cell type in the reference profile are displayed as a heatmap and inform the clustering of rows. **d.** UMAP of 1,862 immune cells colored by cell state type. Inset, UMAP of all cells with immune cluster indicated by red box. **e.** UMAP of immune cells indicating presence of *AHR* expression. **f.** Average immune infiltrate detected in GeoMx^®^ DSP compared to CosMx^®^ SMI analysis for GBM-1. All T cell populations in DSP analysis were combined into a single population to mirror the SMI populations. **g.** Spatial gene expression of *PTPRC* mRNA with inset of immune infiltrate proportions; and Voronoi plots of monocyte, macrophage and microglia in GBM-1 FOV2, 4 and 10. Scale, 100 µm.

To investigate the direct interactions that may be driving immune cells to co-occur more frequently within different tumor niches, we performed ligand/receptor analysis on the whole transcriptomics data from the GBM samples (DSP study). Filtering the ligand/receptor pair interaction based on pattern of expression (ie. tumor to immune, tumor to tumor), we identified interactions that preferentially occurred between tumor and immune cells, such as TGFβ signaling (**Fig. 5a, Supplementary Fig. 11a**). Analysis of a cohort of tumor to immune genes revealed patterns of communication, such as signaling via the TGFβ pathway, where regions enriched for *TGFBR1* signaling expression saw significantly increased BDM and microglia infiltrate (correlation: 0.88, p-value < 0.001), and *ACVR1B* receptor expression correlated with neutrophil and lymphoid infiltrate (correlation: 0.58) (**Fig. 5b**). To gain a better resolution of the individual immune cell types and tumor states involved in these signaling pathways, we applied the top filtered ligand/receptor genes to the SMI data in GBM-1 (**Fig. 5c, Supplementary Fig. 11b**). Strikingly, signaling pathways appeared to dominate the interactions between each tumor state and the immune compartment within. Astrocytic cells were characterized by TGFβ and BMP signaling pathway signaling (*BMP7, TGFB2* expression, **Fig. 5b-e**), supporting findings that increased microglia and BDM infiltrate identified in the DSP study may be due to the high proportion of astrocytic cells in the tumor regions. Mesenchymal cell interactions were dominated by the integrin pathway (*SPP1, VEGFA* expression, **Fig. 5c, e**), Mixed cells signaled predominantly through the WNT pathway (*WIF1, NPPC* expression, **Fig. 5c, e**) and the progenitor subset (NPC, OPC, RG) displayed Notch and PDGF signaling (*DLL1, PDGFA* expression, **Fig. 5c, e**). Notably, the *TGFB1* was an important ligand for signaling in the tumor microenvironment, with receptor specificity to *TGFB1* defining interactions with different tumor and immune cell states (**Supplementary Fig. 12**). This suggests that GBM microenvironments are partially shaped by subtype-specific ligand/receptor signaling, controlling interactions with both tumor and immune cells.

**Fig. 5.**
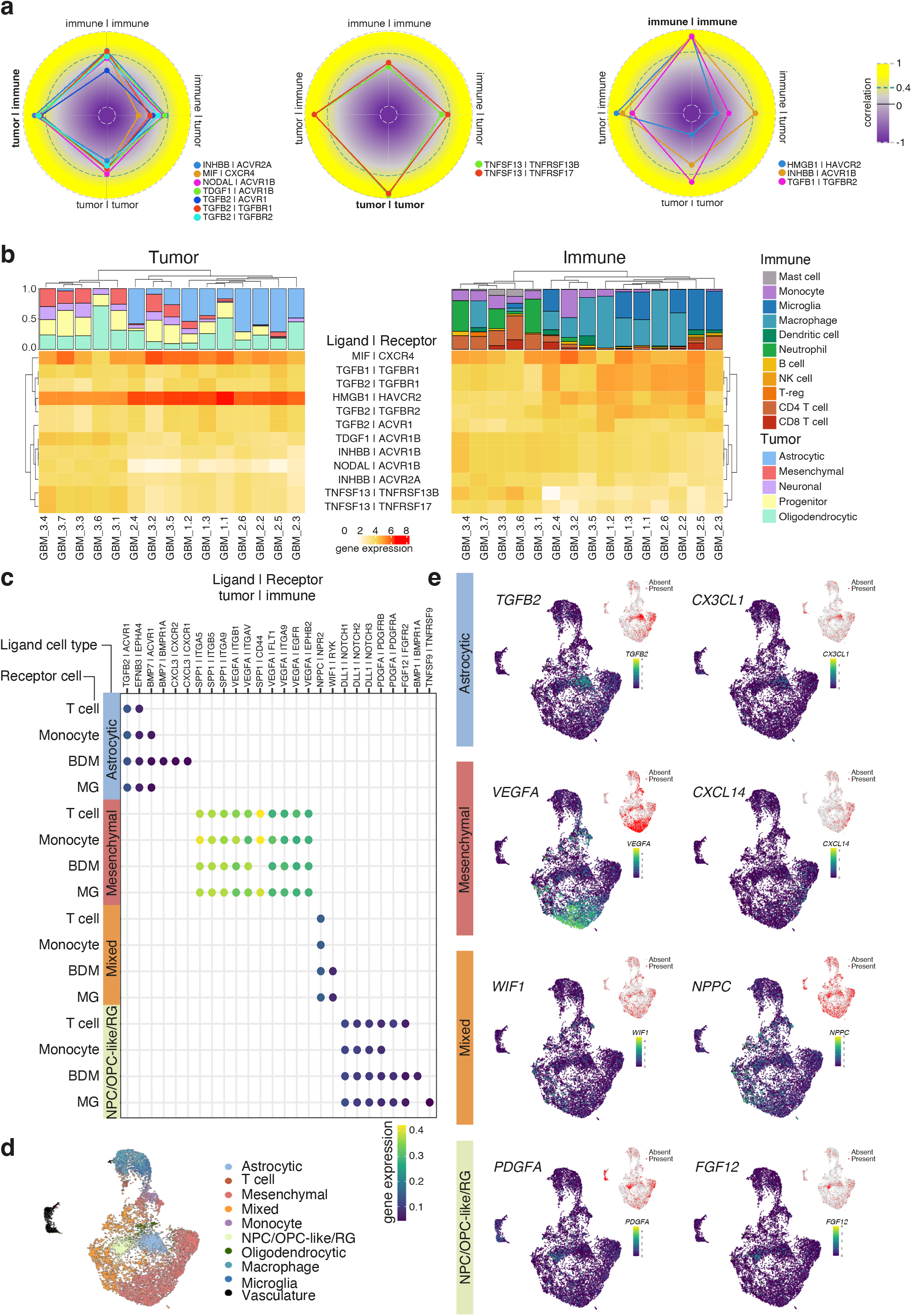
Ligand-Receptor pairs coordinate cells into niches. **a.** Correlation spider plots of ligand receptor pairs with pronounced tumor to immune, tumor to tumor and immune to immune signaling. **b.** Heatmaps of ligand receptor signaling pair expression. Left, cell state deconvolution and heatmap of tumor cell ligand expression. Right, immune deconvolution and heatmap of respective immune cell receptor expression. **c.** Product of expression of significant ligand/receptor interaction pairs between tumor (ligand) and immune (receptor) that show specificity with regards to tumor state in GBM-1. **d.** UMAP of 11,801 cells colored by inferred cell type. **e.** UMAP of all cells showing gene expression level (main) and probe presence/absence (inset) of key genes in each tumor cell state.

To investigate the spatial association of tumor cells within their microenvironment, we identified neighborhoods of similar cell type composition across all FOVs of GBM-1. This approach defined five unique neighborhoods characterized by distinct groups of cells, likely dictated by secreted factors in the microenvironment and receptor/ligand interactions (**Fig. 6a, Supplementary Fig 13a**). The most distinct cellular architecture is observed in neighborhood 4, which is dominated by vasculature (endothelial and mural cells) together with T cells and macrophages, captured entering the tumor via the vessels (**Fig. 6b**). Neighborhood 5 reveals the brain-intrinsic tumor niche (astrocytic, oligodendrocytic and progenitor cell states) together with the microglial cells. This association represents a classical feature of the brain microenvironment under normal conditions, where resident microglial cells associate with glial cells in an absence of lymphocyte infiltration^14,23,24^. In contrast, the mesenchymal and mixed tumor cell populations more broadly associate with circulating immune cells, including macrophages, T cells and monocytes in neighborhoods 1-3 (**Fig. 6b**). To interrogate cell interactions within the neighborhoods, we created network plots and examined proximity between different cell types within approximately 30 µm radius (**Supplementary Fig. 13b**). Neighborhood 1 was dominated by mesenchymal, mixed and monocytes, that cluster into discrete arms highlighting distinct and direct interactions between these cells (**Fig. 6c**). Visualization of these cell states revealed the distinct co-localization in FOV1, and highlighted the clustered nature of Neighborhood 1 (**Fig. 6d, Supplementary Fig. 13c**).

**Fig. 6.**
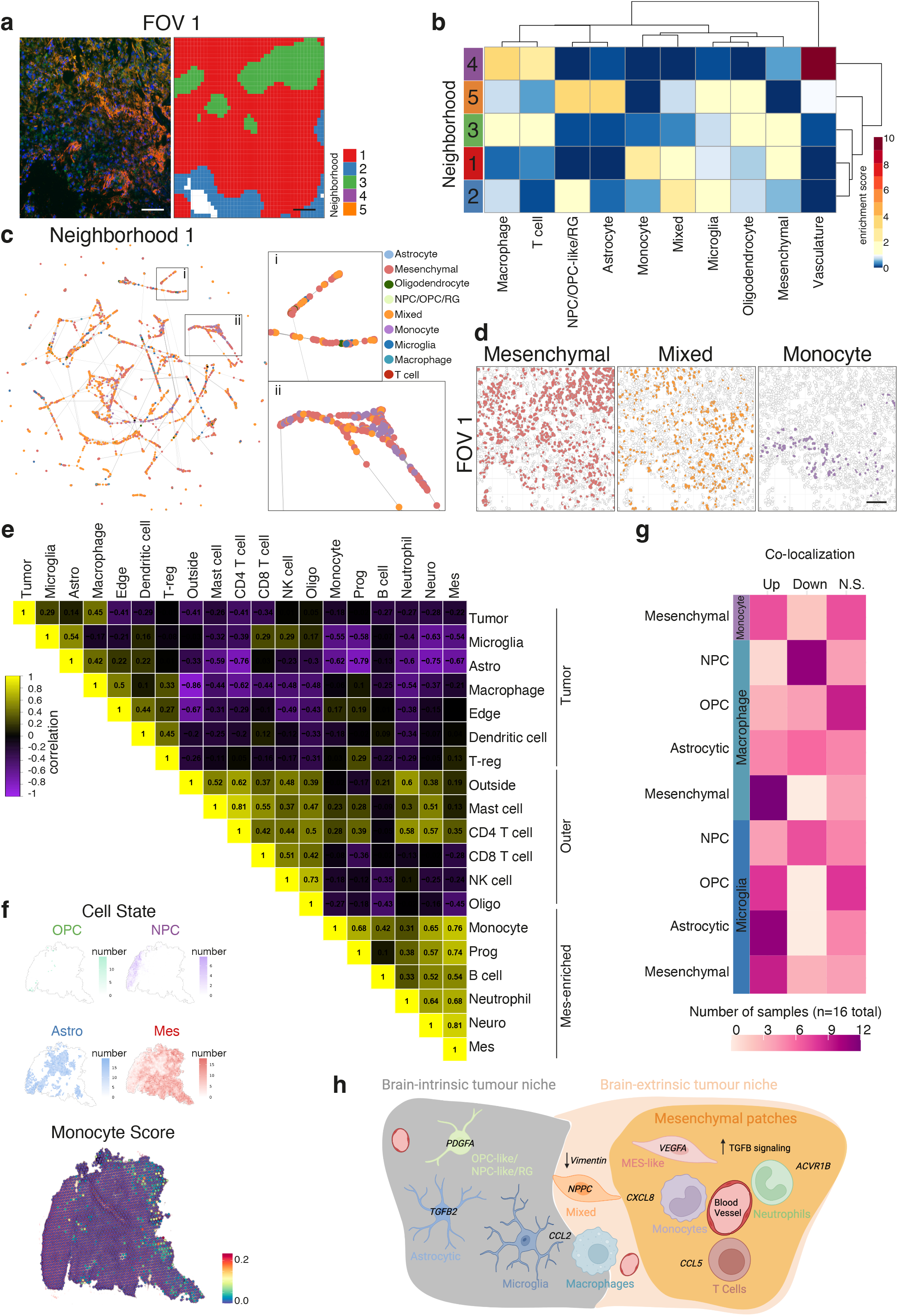
Tumor and immune cell types associate in discrete neighborhoods. **a.** Immunofluorescence image (left) and plot indicating predicted neighborhoods (right) corresponding to spatial niches of field of view (FOV) 1 in GBM-1 SMI analysis. Scale, 100 µm. Enrichment heatmap of each cell type over expected within neighborhoods in GBM-1. **c.** Network where each node represents a cell, colored by their cell type, and each edge represents that another cells is within 30 µm radius for all cells assigned to neighborhood 1. **d.** Voronoi plots of mesenchymal, mix and monocyte populations in FOV 1. **e.** Correlation plot of immune cell types, tumor cell states, and location classifications across matched GBM immune samples (n=15). **f.** Representative GBM Visium^®^ sample^15^ with spots colored by estimated number of cells of mesenchymal, NPC-like, OPC-like and astrocytic tumor state (above) and Monocyte score (below). **g.** Number of Visium^®^ samples^15^ with significant positive or negative co-localization between tumor and immune cell types. **h.** Schematic of tumor and immune architecture in GBM samples highlighting key findings from this study.

To investigate these associations in an independent analysis and across multiple GBM tumors, we next interrogated co-localization of our deconvoluted cell states in the DSP dataset in addition to physical locations in the tumor. Correlation between the cell states identified three domains: tumor, outer and mesenchymal-enriched. Within the tumor domain, astrocytic cells and microglia associate with the central tumor, while at the tumor edge, dendritic cells and regulatory T cells preferentially occur (**Fig. 6e**). Consistent with poor infiltration of lymphocytes, T cells, mast cells and NK cells were enriched in the outer domain, while the mesenchymal tumor cells defined a distinct domain rich in monocytes, neutrophils, B cells and neural progenitor cells (**Supplementary Fig. 14**). Consistent with our SMI analysis of GBM-1, the relationship between mesenchymal cells and monocytes was strong (correlation: 0.76). To extend this observation to a larger independent dataset, we obtained the spatial transcriptomic data from a cohort of GBM (n=16) analyzed in the Visium dataset^14^. Deconvolution of cell types from each spot revealed distinct patches of astrocytic and progenitor cells within the tumor, distinct from the mesenchymal regions (**Fig. 6f**). Consistent with our observations, monocytes localized to the mesenchymal domain in the representative GBM sample and were absent from the regions rich in brain-intrinsic tumor cell types. To extend this analysis across all the 16 GBM samples and broaden to more cell types, we performed co-localization analysis (**Fig. 6g**). We found that microglia most frequently associated with astrocytic tumor cells and macrophages strongly associate with the mesenchymal tumor cells, validating the neighborhood analyses we performed across an independent cohort of patient samples. Taken together, we identify consistent patterns of tumor architecture and immune infiltration across two cohorts of GBM using three independent spatial transcriptomic technologies (**Fig. 6h**). We find a brain-intrinsic tumor niche comprised of astrocytic, oligodendrocytic and neural progenitor tumor cell states associating with the brain resident microglia. A second niche we call the brain-extrinsic tumor niche contains mesenchymal patches infiltrated by monocytes and is rich in TGFβ signaling. Within these patches, neutrophils, T cells, monocytes and macrophages are located near blood vessels and respond to specific ligands via CXCR4. The niche also contains regions marked by the absence of Vimentin where mixed tumor cells comingle with macrophages and monocytes. Collectively, our results provide a model of the GBM microenvironment encompassing discrete niches characterized by the activity of separate signalling pathways.

## DISCUSSION

Here, we provide a spatial characterization of the tumor microenvironment in *IDH* wild type and *IDH1*-mutant high-grade glioma, revealing vast heterogeneity and discrete interactions between tumor cell states and immune cells within a highly organized tumor architecture. Although past studies have identified tumor cell states in glioma and shown the vast heterogeneity of these cell states within individual tumors^13,38,39^, this is the first study to spatially interrogate transcriptional heterogeneity of glioma at a single cell level. Through the adaptation of bioinformatic tools to analyze new datasets of spatial transcriptomics (DSP, SMI), we observe immense molecular and cell composition heterogeneity, which were conserved across both independent cohorts of GBM samples and spatial transcriptomics technologies.

In our study, both GBM and *IDH1*-mutant glioma displayed significant inter- and intra-tumoral clonal heterogeneity as inferred by CNAs. GBM harbors many individual-specific genetic alterations mostly associated with initial malignant transformation, but with minimal evidence for regional variation as observed by multi-site sampling^40^. We also find heterogeneity between patients, but in contrast to previous work, our admittedly limited GBM samples showed significant evidence of intra-tumoral heterogeneity. Our finding closely aligns with work by others utilizing spatial transcriptomics that finds regional accumulation of CNAs in response to hypoxia^15^.

Recent stratification of GBM based on cell lineages of the normal fetal brain characterized five cell states in two broad groups^13^. The normal neural lineages: astrocytic, oligodendrocytic and neuronal, together with the proliferative progenitor cell state; and the mesenchymal lineage which does not have a direct parallel in the normal brain. Like previous studies that rely on multi-site sampling from different anatomical locations^41^, we demonstrate tumors contain multiple subtypes sometimes in close proximity. We extend these initial observations in the context of their spatial relationship with the immune compartment, that defines a brain-intrinsic niche – comprised of astrocytic, oligodendrocytic, neuronal and progenitor cell states – closely associated with the resident microglial cells and rarely interacting with immune cells migrated from the circulation. In contrast, a brain-extrinsic niche characterized by mesenchymal tumor cells, closely associated with macrophages, monocytes and other lymphocytes, is consistent with recently described mesenchymal patches in GBM that are clearly spatially demarcated^14^, and thought to be linked to tumor progression^42^. The nature of immune infiltration in mesenchymal patches concurs with work by *Varn et al.* finding that myeloid cells in mesenchymal tumor subtypes are likely to be macrophages^20^. The mixed tumor state, possibly representing cells differentiating towards the mesenchymal lineage, is preferentially located in brain-extrinsic niches. We also show that the conditions in brain-intrinsic and extrinsic tumor niches are likely dictated by specialized ligand/receptor interactions in agreement with Gangoso *et al.* who demonstrated tumor state specific transcriptional and epigenetic changes reshape the immune microenvironment^21^. The immune and regulatory underpinning of the dichotomy of the glioma microenvironment suggests that the success of immunotherapy agents such as chimeric antigen receptor (CAR) T cell infusion, checkpoint inhibitor blockade or dendritic cell therapies^43^, may be influenced by tumor composition.

In *IDH*-mutant glioma, scRNAseq analysis had identified the presence of cell states restricted to the astrocytic and oligodendrocytic lineages^16,17^. Here, we demonstrate the dominance of the oligodendrocytic lineage across both astrocytoma and oligodendroglioma samples, conserved spatially throughout the tissue. The neuronal lineage was also present in these tumors and increased in frequency as the tumor purity declined, likely representative of normal neural cells. Notably, progenitor cells were identified in the tumor regions, and stratified to the proliferative tumor compartment only, consistent with previous reports of a rare subpopulation of progenitor cells^16^.

Indeed, the intra-tumoral heterogeneity of *IDH1-*mutant high-grade glioma was substantially reduced in comparison to GBM. In contrast, inter-tumoral heterogeneity was increased, though this was driven by the oligodendroglioma re-diagnosis of A-1, which limited our ability to closely examine the *IDH1*-mutant cohort. This marked difference in heterogeneity between grade 3 and grade 4 gliomas has been postulated to reflect a more lineage-restricted cell-of-origin in *IDH-* mutated tumors^17^.

Heterogeneity is a hallmark of GBM and has been on display in this spatial analysis in comparison to astrocytoma, as in other recent GBM datasets^14,15^, with implications for the limitations of personalized therapies for patients. Further, spatial heterogeneity highlights the challenge of identifying biomarkers for treatment from potentially un-representative diagnostic sampling, and will likely create a hurdle in future clinical decision making. We, and others, have highlighted the association of the mesenchymal lineage with increased infiltration of circulating immune cells and as described above, may represent an avenue to leverage immunotherapeutic approaches, in patients who are more likely to respond. This is in contrast to chemotherapy treatment, whereby mesenchymal cells appear to be resistant to standard therapy and are frequently observed in recurrent tumors^19,42^, requiring novel treatment strategies. Thus, routine mechanisms to predict the proportion of mesenchymal lineage cells would likely aid in diagnostic and prognostic predictions. We show that the combination of central tumor (CT) annotation with the proliferative state (Ki67 immunostaining) may provide meaningful information in estimating the prevalence of progenitor and mesenchymal cell lineages in the tumor, which could aid assessment of the propensity of the tumor to recur and provide insight into the cell state of the GBM in the absence of advanced genomic testing. Future studies employing high-plex spatial molecular imaging with an increased sample size will aid in the further characterization of spatially distinct domains to identify mechanisms of plasticity and immune infiltration and their secreted factors in patients with high-grade glioma. Ultimately, understanding the spatial distribution of tumor cells within their microenvironment will provide clinical impact with the potential to selectively modulate cellular plasticity and reshape the landscape of therapeutic targeting in GBM.

## METHODS

### Clinical Samples

All tissue samples were obtained from the Royal Melbourne Hospital Neurosurgery Brain and Spine Tumour Tissue Bank (Melbourne Health Ethics # 2020.214). Formalin fixed paraffin embedded (FFPE) tissue were sectioned at 5 μm thickness and mounted onto Superfrost Plus slides within one month of spatial transcriptomics analysis. Slides were stored at 4 °C with desiccator and shipped to NanoString (Seattle, USA) for Technology Access Program (TAP) processing for both GeoMx^®^ DSP and CosMx^®^ SMI analyses.

Immunohistochemistry of GFAP (Dako Z0334; 1:50 dilution; epitope retrieval: Roche Ventana ULTRA Cell Conditioning Solution 1 pH 8.0-8.5 (CC1) 32 minutes at 36 °C), IDH1-R132H (Dianova IDA-H09; 1:50 dilution; epitope retrieval: CC1 32 minutes at 100 °C), CD45 (Dako M0701; 1:1000 dilution; epitope retrieval CC1 32 minutes at 100 °C), KI67 (Dako M7240; 1:100 dilution; epitope retrieval: CC1 32 minutes at 100 °C) and PDGFRA (Abcam ab203491; 1:200 dilution; epitope retrieval: EDTA buffer, pH 8.0-8.5 at 100 °C for 15 minutes) was performed. Hematoxylin and eosin (H&E) stained sections and immunohistochemistry slides were scanned using the 3D Histech Brightfield (20X) and processed using CaseCentre online software.

Pathology review was performed on all the six cases. The tissue sections demonstrated distinct morphological features and were assessed using the cellularity of the tumor cells, nuclear pleomorphisms, presence of mitoses, necrosis, and microvascular proliferation. The five anatomic features proposed by The Ivy Glioblastoma Atlas Project (Ivy GAP) were used to annotate the tumors into regions. These included: Leading edge (LE): the outermost part of tumour with the least tumor cell density and bordering on the normal tissue area; Cellular tumor (CT): the core of the tumor with highest cell density; Infiltrating tumor (IT): the zone interposed between LE and CT with moderate tumour cell density; Microvascular proliferation (MVP): Areas with endothelial proliferation; Pseudopalisading cells around necrosis (PAN): dense tumour cells geographically fenced around core necrosis. Other features recognized are regions of large non-palisaded necrosis (NEC) and normal brain tissue (N). Annotations were performed by two pathologists.

The evaluation of Ki67 and CD45 immunostaining was performed according to standard established protocols using light microscopy. The intensity and percentage of tumour Ki67 positive cells were assessed semi-quantitatively. The intensity was graded as 0, +, ++ and +++ representing negative, weak, moderate, and strong staining respectively. The percentage was calculated by dividing the number of positive cells (minimum number required was 200 positive cells) by total number of tumour cells in the tumour areas assessed as a percentage. Multiple areas of tumours were counted and the average range of percentages were recorded for each case. The intensity of CD45 immunostaining was semi-quantitatively graded as 0, +, ++ and +++ representing negative, weak, moderate, and strong staining respectively. Each tumour was assigned the grade corresponding to its degree of immunoreactivity.

### GeoMx**^®^** Dataset

#### Digital Spatial Profiling (DSP)

After baking the slides for 2 hours for paraffin removal, slides were loaded onto a Leica BOND RX for tissue rehydration, heat-induced epitope retrieval (ER2 for 20 minutes at 100 °C) and proteinase K treatment (0.1 μg/ml for 15 minutes at 37 °C). The tissue sections were then hybridized with the Human WTA probes overnight. Following 2 x 5 min stringent washes (1:1 4x SSC buffer & formamide), the slides were blocked and then incubated morphology marker antibodies: GFAP (488 channel, NBP2-3318AF488, Novus), CD45 (594 channel, 13917BF, Cell Signaling Technologies), and Ki67 (647 channel, 9027BF, Cell Signaling Technologies). Syto83 (532 channel, S11364, Invitrogen) was used as a nuclear stain. Tissue sections were then loaded into the GeoMx^®^ platform and Regions of interest (ROIs) were selected guided by the immunofluorescence of the morphology markers antibodies listed above. Segmentation was used in some ROIs used to enrich for Ki67+ and/or CD45+ populations. UV light was then directed by the GeoMx^®^ at each AOI and released the RNA ID and UMI-containing oligonucleotide tags from the WTA probes for collection and sequencing preparation. Illumina i5 and i7 dual indexing primers were added to the oligonucleotide tags during PCR to uniquely index each AOI. AMPure XP beads (Beckman Coulter) were used for PCR purification. Library concentration as measured using a Qubit fluorometer (Thermo Fisher Scientific) and quality was assessed using a Bioanalyzer (Agilent). Sequencing was performed on an Illumina NovaSeq 6000 and .fastq files were processed into gene count data for each AOI using the GeoMx^®^ NGS Pipeline.

#### Preprocessing

For all computational analysis methods were run using defaults unless otherwise specified. Raw counts from the two separate experimental batches were grouped into a single dataset. AOIs were removed if their nuclei count was below 100 nuclei, as provided by NanoString. Genes were filtered out unless above limit of quantitation in 5 % of remaining AOIs, calculated for each AOI as the geometric mean of all negative probes multiplied by the geometric standard deviations of the negative probes to the second power.

From filtered raw counts of each region, a combined dataset was constructed, where Ki67 (K) and Ki67^-^ (T) tumor AOIs from the same region were joined together into a single tumor AOI ([T + K]). This was done to ensure the entire neoplastic compartment of each region was being compared when desired. Further preprocessing of both the original separated [T] + [K] dataset and the combined [T + K] dataset were performed independently but identically as follows.

Remaining genes were normalized by upper quartile normalization. Batch correction was required as we observed different signal-to-noise ratios, as previously observed^44^. Here we performed batch correction on the logged normalized count data, using limma via geomxBatchCorrection in standR (v1.0.0), separately for astrocytoma and GBM, with experimental batch and sex considered. Correcting for batch effects on all astrocytoma and GBM samples simultaneously showed inappropriate mixing of tumor classes in PCA and suggested loss of biologically relevant differences. PCAs were plotted using functions from the standR package (v1.0.0). Cell cycle scoring was estimated by Seurat (v4.1.1). Hypoxia levels were measured by abundance of genes in each AOI from the respective KEGG pathway using the keggGet function (KEGGREST v1.36.3).

#### Deconvolution

The tumor deconvolution reference was constructed from Couturier *et al.* dataset with unassigned tumor cells excluded^13^. The top 5,000 highly variable genes also present in the DSP filtered gene list were used to form signature profiles for each cell state, via the spatialDecon package (v1.6.0). Testing for difference in cell state proportions between astrocytoma and GBM as well as Ki67^+^ and Ki67^-^ regions was performed using propeller in the speckle package (v0.0.3), using t-tests and estimated number of nuclei of each cell state. A logit transformation was used to test the proportions and multiple testing was corrected using false discovery rate (FDR).

The immune deconvolution reference was constructed from Ruiz-Moreno *et al.* dataset^14^ via the spatialDecon package (v1.6.0). Immune cell type signatures were derived from Ruiz-Moreno ‘cell type’ annotation; for segregating T cells, only cells that were labelled as ‘mature T cell’ in the ‘cell_type’ annotations and contained either ‘CD4’, ‘CD8’, or ‘reg’ in the ‘celltype_original’ annotations were included for each respective cell type. Alongside the immune cell types, a neoplastic signature and non-immune non-neoplastic signature were also included from ‘annotation_level_1,’ to account for possible contamination of immune AOIs. All genes intersecting the safeTME profile as well as the DSP filtered gene list were included (n = 859). Testing for differences in cell type proportions between classifications was performed using propeller in the speckle package (v0.0.3), using F-tests via ANOVA and estimated number of nuclei of each cell state. A logit transformation was used to test the proportions and FDR was applied.

#### DEG

DEG analysis was performed with limma-voom (limma v3.52.4) /edgeR (v3.38.4). Voom was applied twice, with duplicate correlation considered both times for AOIs coming from the same region. Sex and Batch were also added to the model where appropriate. GO term analysis was performed on DEG lists using goana, with trend set to TRUE (limma v3.52.4). GO terms were then clustered using semantic clustering via simplifyEnrichment (v1.6.1), and further clustered with kappa clustering to separate into groups of similarly worded GO terms that share similar gene sets.

#### CNA analysis

CNA analysis was performed using cnvkit (v0.9.9) to infer copy number alterations from raw count data. GBM samples from The Cancer Genome Atlas with matched CNA and gene expression data were used to calculate the correlation of each gene’s expression to copy number changes. Immune AOI were used as ground truth normal samples to which alterations were compared to. Circular binary segmentation was performed. CNA results were visualized using ComplexHeatmap (v2.12.1) and GenomicRanges (v1.48.0), with windows binned to 10,000 base pairs using EnrichedHeatmap (v1.26.0).

#### Shannon Entropy

Shannon entropy across all individual genes was calculated from upper-quartile normalized counts using a shrinkage estimator to counter the effect of under sampling for lower nuclei AOIs. For entropy of tumor cell states, the entropy of the top 50 scaled genes contributing to the respective cell state signature in the Couturier *et al.* reference profile was summed together^13^. Entropy was calculated for various subsets of the data, including: the combined [T + K] tumor samples; the separated [T] + [K] tumor samples; the Ki67^+^ tumor regions only; and the Ki67^-^ tumor regions only.

#### Sum of Squares approach

A sum of squares approach was applied to each window of 10,000 base pairs along the genome. For every chromosomal window assessed, the total sum of squares is the combined result of: the between sum of squares, which aggregates the difference of each sample’s average value to the global average (inter-tumoral heterogeneity); and the within sum of squares, which aggregates the difference of each AOI of a sample to that sample average, and then combines values for all samples (intra-tumoral heterogeneity). This approach was applied separately to the GBM samples and the astrocytoma samples, revealing genomic regions of heterogeneity for each glioma.

#### Correlation plot

GBM ROIs with matched immune and tumor AOIs were correlated along with their location classifications based on immunostaining, employing the combined [T + K] dataset. ROIs labelled as ‘Ki67 Border’ and ‘Normal regions’ constituted the ‘Outside’ locations, and ROIs labelled as ‘Border’ formed the ‘Edge’ location signature. The correlation plot was made using corrplot (v0.92).

#### Ligand-receptor analysis

Ligand-receptor analysis employed the connectomeDB2020 list of literature-supported ligand-receptor pairs. Only ROIs with matching tumour and immune AOIs were included. Correlations between ligand and receptor expression was calculated in the same AOI (immune to immune, and tumor to tumor signaling) and in matched AOI pairs (tumor to immune, and immune to tumor signaling). Displayed ligand-receptor pairs were manually selected based on the recurrent increased correlation of particular signaling pathways in the same direction (for example, increased correlation of WNT signaling pairs in tumor to tumor interactions). Spiderplots were constructed with ggradar (v0.2).

### CosMx**^®^** Dataset

#### Spatial Molecular Imaging (SMI)

Tissue samples were obtained and prepared for analysis as previously described^27^. Briefly, serial sections were H&E stained to identify FOVs for RNA target readout and protein detection. RNA readout was performed by flowing 100 μl of Reporter Pool 1 into the flow cell and incubating for 15 min. Reporter Wash Buffer (1 mL) was then flowed to wash unbound reporter probes, and Imaging Buffer was added for imaging. Nine Z-stack images (0.8 μm step size) for each FOV were acquired, and photocleavable linkers on the fluorophores of the reporter probes were released by UV illumination and washed with Strip Wash buffer. The fluidic and imaging procedure was repeated for the 16 reporter pools, and the 16 rounds of reporter hybridization-imaging were repeated multiple times to increase RNA detection sensitivity. After RNA readout, the tissue samples were incubated with a 4-fluorophore-conjugated antibody cocktail against Histone H3, 18s rRNA, GFAP and DAPI stain in the CosMx SMI instrument for 2 h. Imaging Buffer was added to the flow cell and nine Z-stack images for the 4 channels (3 antibodies and DAPI) were captured.

#### Cell segmentation

For cell segmentation the NanoString pipeline combining image preprocessing and machine learning techniques was used on tissue images stained with Histone H3, 18s rRNA, GFAP and DAPI. Briefly, the pipeline first performs pre-processing for boundary enhancement followed by cell segmentation via a pretrained neural network in Cellpose (v2.0).

#### Preprocessing

We next generated single-cell expression matrix by counting molecules of each gene within the area assigned to a cell by the segmentation algorithm. We removed any cells with fewer than 20 total transcripts or more than 3 negative control probes. We then applied a log-normalization to the expression matrix using scater (v1.26.1) in combination with SpatialExperiment (v1.9.4). Highly expressed *MALAT1* was removed before any subsequent analyses.

#### Cell type annotation

Data dimensions were reduced using PCA and the number of components kept was determined via a global maximum likelihood based on translated Poisson mixture model approach with 20 nearest neighbors implemented in intrinsicDimension (v1.2.0). This approach has recently been shown to produce the best separation of challenging subpopulations.We also generated a Uniform Manifold Approximation and Projection (UMAP) embedding based on the PCA with *spread* set to 3 and *mindist* set to 0.01 with 15 nearest neighbors. Cells were then clustered based on the PCA via Leiden clustering implemented in bluster (v1.8.0). Cells were automatically annotated using SingleR (v2.0.0) with the reference set as the downsampled GBM harmonized dataset^14^ (see description in Deconvolution).

Clustering of the entire dataset produced 35 clusters. Clusters were first annotated as Vasculature, Immune or Tumor according to the predicted cell type labels from SingleR and marker genes (Vasculature: *COL4A2*, Tumor: *APO-J*, Immune: *CD163*). Cells in the immune and tumor compartment were reclustered following the process outlined above, including dimension reductions (during the immune cell reclustering we identified further mesenchymal cells). This produced 16 clusters for the immune cells and 27 clusters for the tumor cells. Cell type labels for the tumor cells were additionally predicted using SingleR with the reference set to the Couturier *et al.* dataset^13^. Clusters for both immune and tumor cells were annotated according to the predicted labels and known markers genes.

#### Networks and modularity

Using reticulate (v1.28), the preprocessed dataset was converted to an *anndata* object to be analyzed by squid.py (v1.2.3). We were then able to build an adjacency matrix indicating the cells that are in ∼30µm radius (179 pixels) of each cell within the same FOV. By combining the adjacency matrix with the cell type annotations for each cell the modularity of each cell type can be worked out using igraph (v1.4.0). The modularity measures how strongly separated cells of the same type are compared to a random null model with higher values showing less spatial separation between cells of the same cell type.

#### Neighborhood clustering

To identify tumor niches, we developed our own version of a neighborhood composition approach. Briefly, we first divide each FOV into overlapping 500 by 500 pixel windows that are shifted by 100 pixels. In each window, we count the number of different cell types. Using a Bray-Curtis dissimilarity implemented in vegan (v2.6-4), we built a hierarchical clustering tree utilizing *hclust.* With the help of the hierarchical clustering tree, we decided on 5 clusters and identified cluster labels for each window using *cutree*. As each 100 by 100 pixel tile of each FOV is overlapped by multiple windows with an associated cluster label, we use a majority voting strategy to identify the cluster label for each 100 by 100 pixel tile. This results in neighborhoods being identified across the FOVs, which can be described by their enrichment of cell types using a simple chi-square test approach.

#### Ligand-receptor analysis

Based on the adjacency matrix, we identified significant ligand-receptor interactions (adjusted p-value <0.01) according to the CellPhoneDB algorithm as implemented in squid.py. We selected ligand-receptor interactions that were previously found to occur mainly in one direction; either tumor to tumor, tumor to immune, immune to tumor or immune to immune, such as the genes in the TGFB pathway. We also selected ligand-receptor interactions that were specific to the ligand cell type.

### Visium^®^ Dataset

Processed data were downloaded from https://doi.org/10.5061/dryad.h70rxwdmj. Deconvolution of spots as described in Ravi *et al.* were obtained from the authors upon request^15^.

#### Co-location testing

To test co-location of two cell types, we modified a method for the identification of proximal interacting cell types in Giotto. We first determined whether the cell types were present in any spot by multiplying the predicted proportion with the number of cells in the spot. Like for the SMI data, we built an adjacency matrix indicating the direct neighbors for each spot. Using this adjacency matrix, we can then find the number of times the cell types of interest are co-located. By permuting the cell type labels 1000 times, we can find distribution of the number of expected co-location events, which allows us to determine a z-score and can be tested. P-values from tests were corrected for multiple testing using *p.adjust* with a FDR method.

### Data availability

The GeoMx^®^ DSP data and CosMx^®^ SMI is available from the authors on request during the review period and will be made available in public repositories upon publication.

### Code availability

All code required for the analysis of the data can be found at: https://github.com/SaskiaFreytag/spatial_brain_cancer/.

## Supporting information

Supplementary package

## ACKNOWLEDGEMENTS

We thank S. Stylli, K. Drummond and J. Dimou for expert curation of the Royal Melbourne Hospital Neurosurgery Brain and Spine Tumour Tissue Bank, M. Bisignano for assistance at the RMH Anatomical Pathology Department and E. Tsui for assistance at the WEHI Histology core. We thank D. Merino and M.L. Asselin-Labat for critical review of the manuscript. This work was supported by a Technology Access Program (TAP) grant to S.A.B. for GeoMx® DSP analysis of two samples, supported by A. Venschoiack, R. John, M. Feterl and B. Bassam. S.A.B. is supported by a Victorian Cancer Agency Mid-Career Research Fellowship (MCRF22003), J.J.D.M. is supported by a WEHI Johnson PhD Scholarship and an Australian Government Research Training Program (RTP) Scholarship, O.E.F. is supported by a WEHI IPSI Scholarship and S.F. is supported by a National Health and Medical Research Council of Australia (NHMRC) Ideas Grant (GNT1184421). This work was made possible and financially supported in part through the authors’ membership of the Brain Cancer Centre, support from Carrie’s Beanies 4 Brain Cancer, a Priority-Driven Collaborative Cancer Research Scheme Grant funded by Cancer Australia to S.A.B. (2003127), and through Victorian State Government Operational Infrastructure Support and Australian Government NHMRC Independent Research Institutes Infrastructure Support Scheme (IRIISS).

## AUTHOR CONTRIBUTIONS

S.A.B., S.F. and J.R.W. conceived the study. J.J. and A.M. collected patient tissue and A.P., D.S., and L.Z. processed the sequencing in the NanoString technology access program. J.J.D.M., L.F. and S.F. performed the bioinformatic analysis. O.E.F. and S.R.T. performed the pathology analysis. S.A.B., S.F., J.R.W., and J.J.D.M wrote and revised the manuscript with input from all authors.

## COMPETING INTERESTS

S.A.B. received instrument support (GeoMx®) from NanoString Technologies as highlighted in the Acknowledgments section. A.P., D.S., L.Z. and Y.L. are employees and stockholders of NanoString Technologies.

## Notes

https://doi.org/10.5061/dryad.h70rxwdmj

### Summary of Updates

Minor modifications to text and figures, including increased statistical testing in the text and alternative layout in Figure 5.

