## Supplementary package for "Spatial architecture of high-grade glioma reveals tumor heterogeneity within distinct domains"

**Author's Affiliations:**

[1]Personalised Oncology Division, The Walter and Eliza Hall Institute of Medical Research, Melbourne, Australia

[2]Department of Medical Biology, University of Melbourne, Melbourne, Australia

[3]Department of Surgery, Royal Melbourne Hospital, Melbourne, Australia

[4]Department of Anatomical Pathology, Royal Melbourne Hospital, Melbourne, Australia

[5]NanoString Technologies Inc., Seattle, WA, USA

[6]Department of Medical Oncology, Peter MacCallum Cancer Centre, Melbourne, Australia

*** Co-senior and corresponding Authors:**

Sarah A. Best,; Saskia Freytag,; James R. Whittle,; The Walter and Eliza Hall Institute of Medical Research, Parkville, Victoria, Australia, 3052. Phone: +61-3-9345-2677; Fax: +61-3-9345-0852.

**Keywords:** Brain Cancer, Glioma, Spatial Transcriptomics, Sequencing, Deconvolution, Immune microenvironment

### Supplementary Information

### Supplementary figure 1

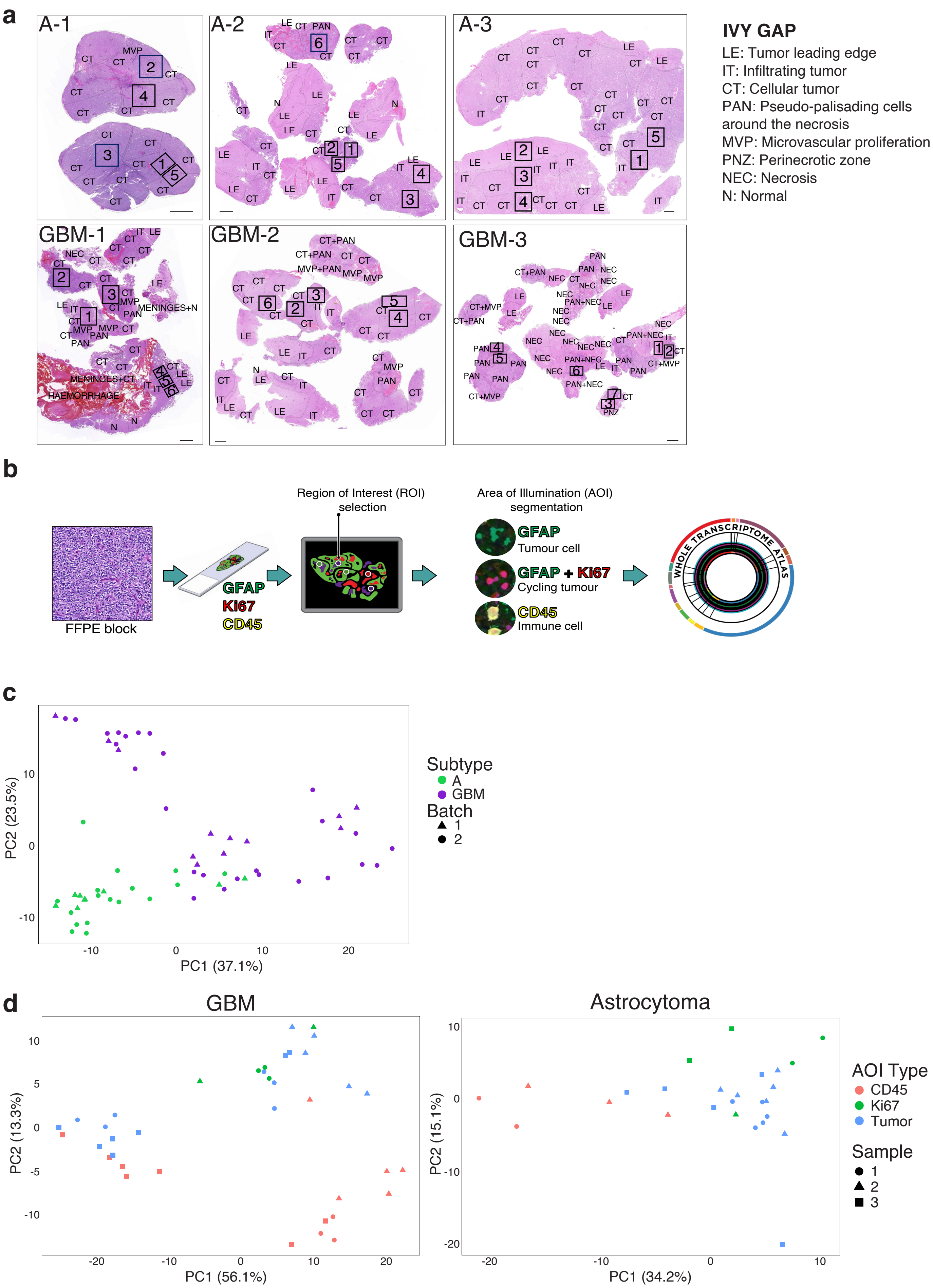

**Supplementary Fig. 1 |**

**a.** H&E of samples with pathology annotation and regions of interest (ROI) indicated. Scale, 1 mm. **b.** Schematic of DSP GeoMx® whole transcriptome analysis workflow. **c.** Principal component analysis of all samples. Displaying first two principal components for all AOIs following batch correction, colored by tumor type. **d.** Principal component analysis of GBM (left) and astrocytoma (right) samples. Displaying first 2 principal components following batch correction, colored by area of illumination (AOI).

### Supplementary figure 2

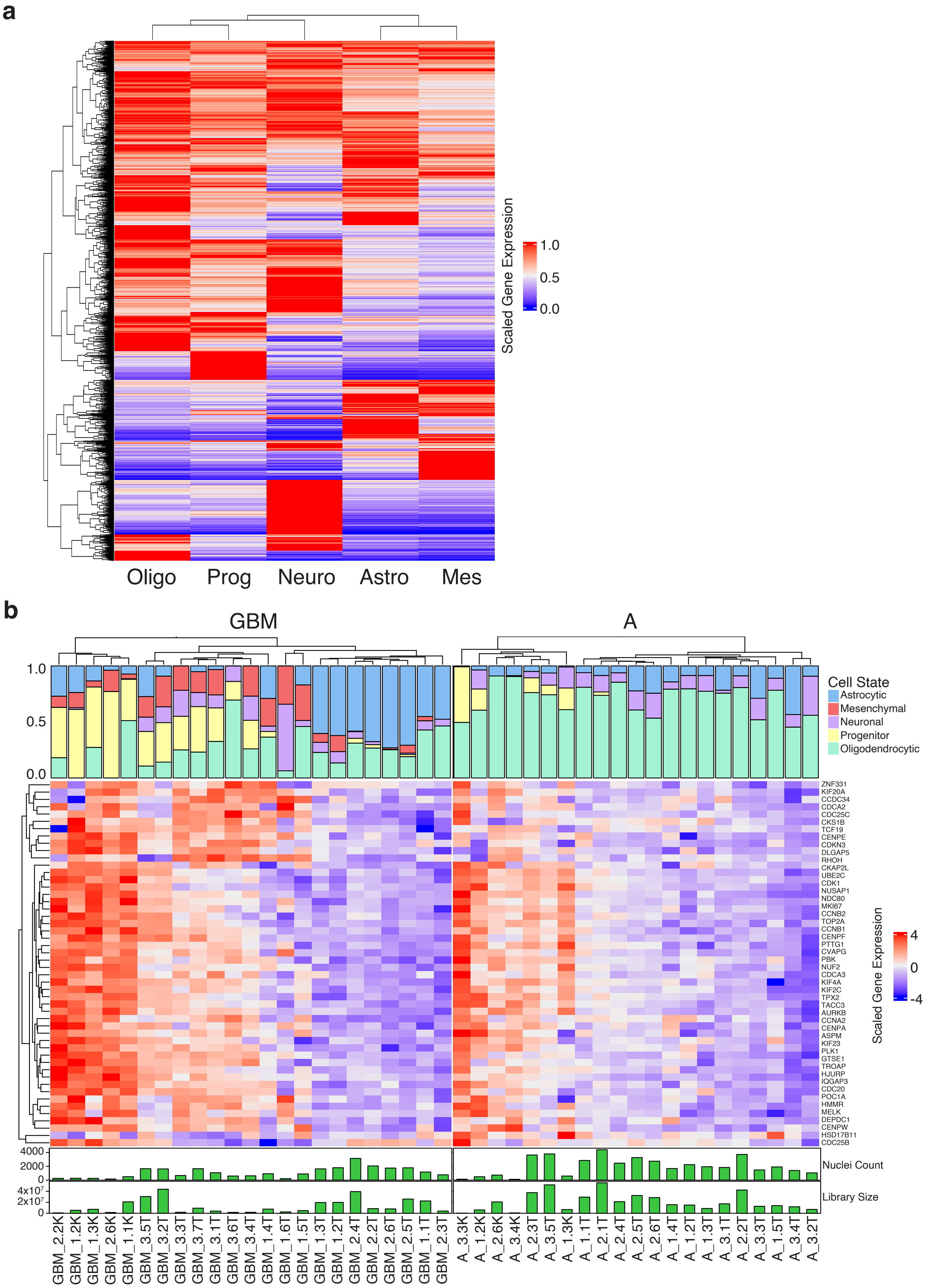

**Supplementary Fig. 2 |**

**a.** Deconvolution heatmap signature. Scaled gene expression of top 5,000 highly variables genes from Couturier *et al.* dataset present in preprocessed gene list, for each cell state profile. **b.** Complex heatmap with cell state and progenitor signature gene expression. Columns clustered by scaled expression of top 50 genes contributing to progenitor signature, annotated below with nuclei count and library size for each AOI.

.

### Supplementary figure 3

**a**

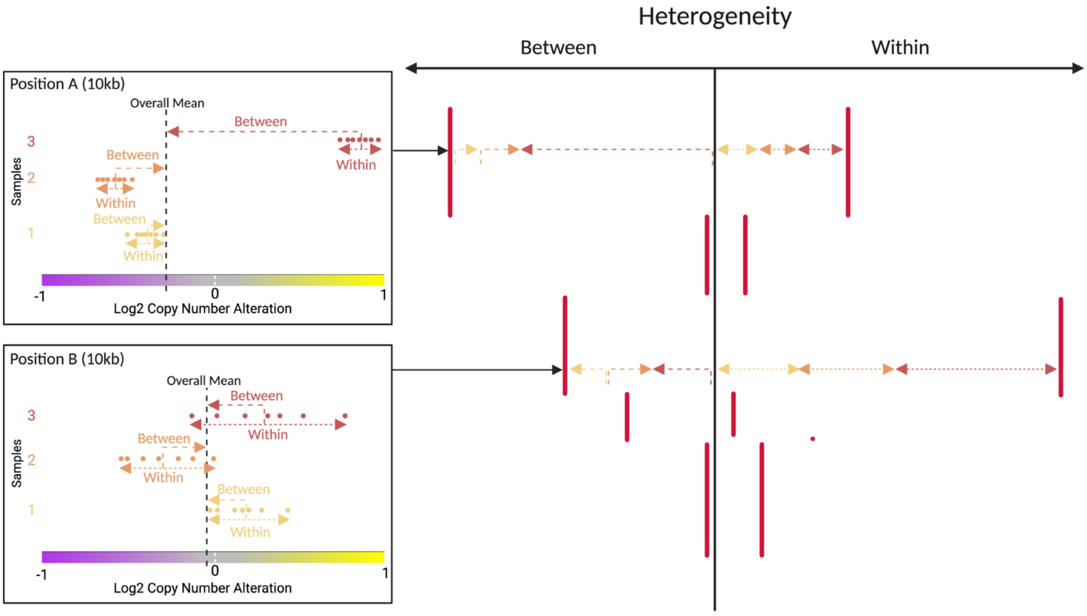


**b**

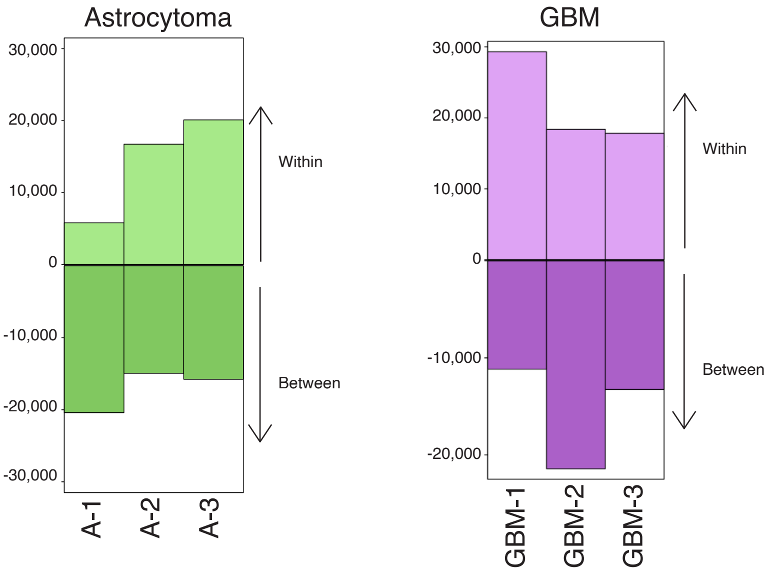

**Supplementary Fig. 3 |**

**a.** Representative heterogeneity schematic outlining the calculation of between and within scores for each sample, and as a whole. For every 10 kb chromosomal window assessed, the total sum of squares is the combined result of: the between sum of squares, which aggregates the difference of each sample's average value to the global average; and the within sum of squares, which aggregates the difference of each AOI of a sample to that sample average, and then combines values for all samples. Created with biorender.com **b.** Between and within calculations for astrocytoma (left) and GBM (right) samples.

### Supplementary figure 4

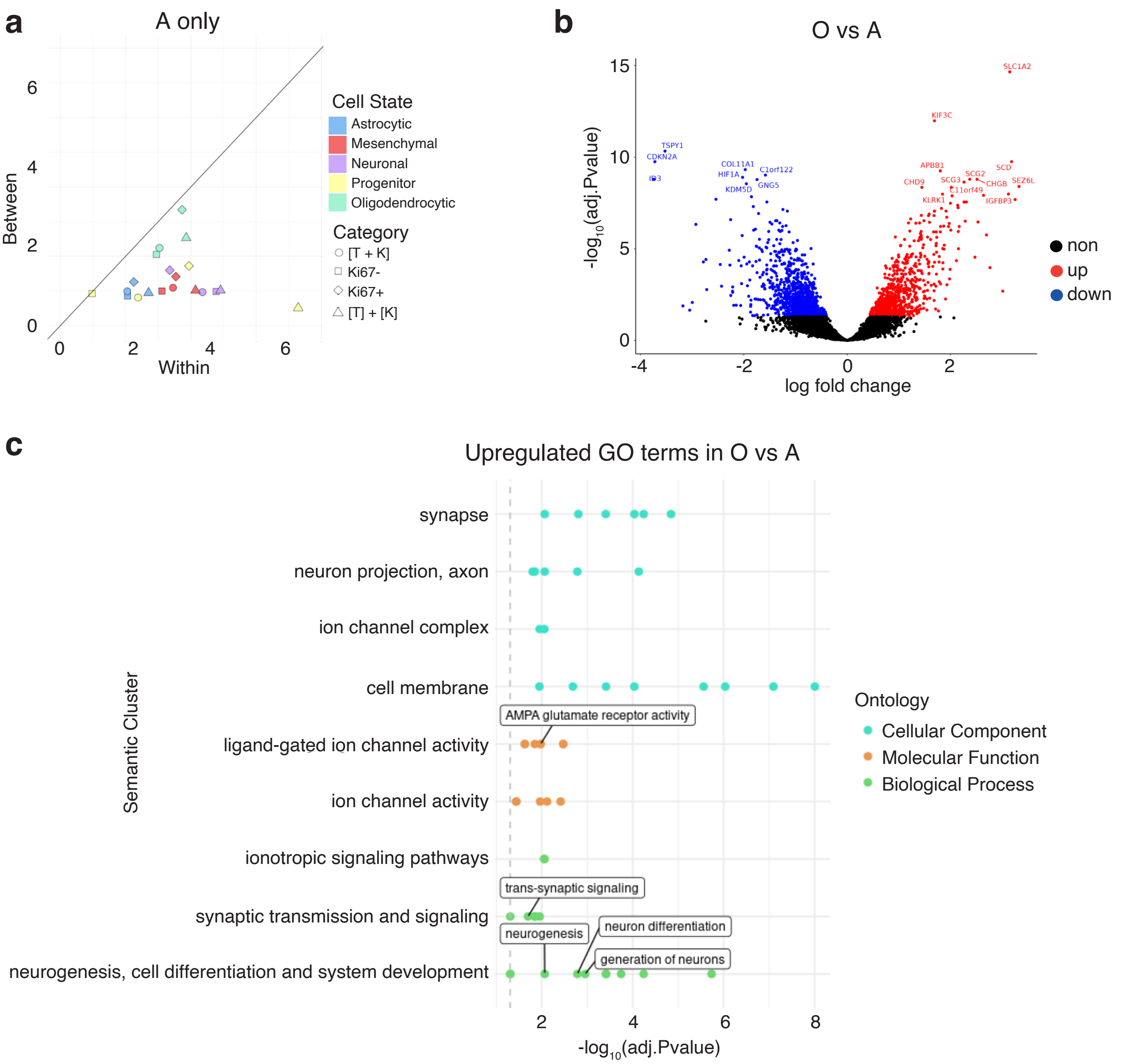

**Supplementary Fig. 4 |**

**a.** Shannon entropy plot of cell state astrocytoma tumour samples only (A-2 and A-3). Colored by cell state, and repeated for both combined ( [T + K] ) and separated ( [T] + [K] ) datasets, as well as $Ki67^-$ and $Ki67^+$ samples only (**Methods**). **b.** Volcano plot of gene expression in astrocytoma (A-2 and A-3) vs oligodendroglioma (A-1) samples. Genes labelled ‘up’ (red) have significantly higher expression in the oligodendroglioma sample, whereas genes labelled ‘down’ (blue) have significantly higher expression in the astrocytoma samples. **c.** Summary of upregulated GO terms in oligodendroglioma (O, n = 1) vs astrocytoma (A, n = 2). GO terms grouped by semantic clustering. Ten of the most numerous and interesting groups are displayed, with key terms highlighted.

### Supplementary figure 5

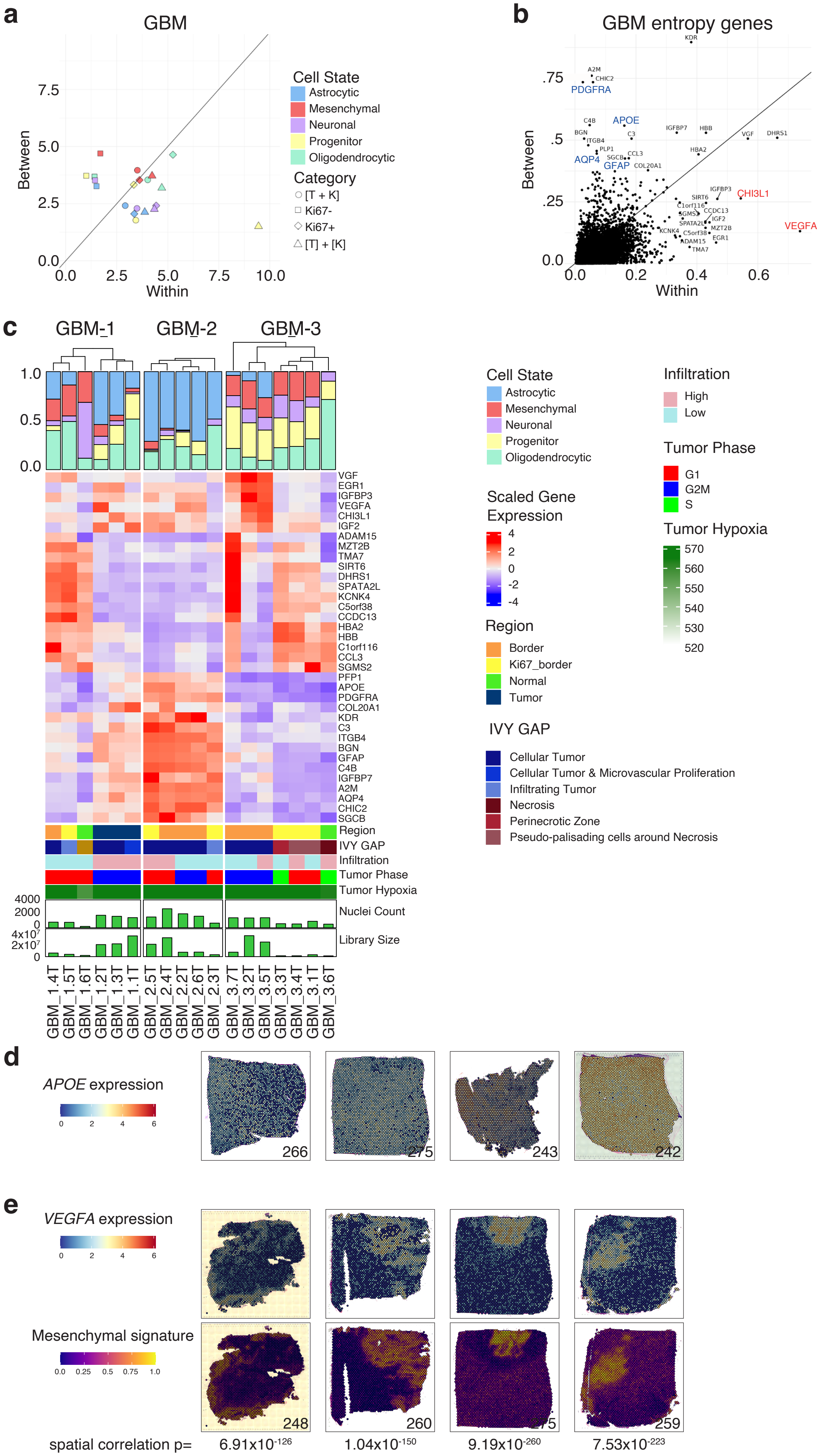

**Supplementary Fig. 5 |**

**a.** Shannon entropy analysis of cell state in GBM tumour samples. Colored by tumor state, and repeated for both combined ( [T + K] ) and separated ( [T] + [K] ) datasets, as well as Ki67$^{-}$ and Ki67$^{+}$ samples only (**Methods**). **b.** Shannon entropy analysis of all genes in GBM tumor samples. **c.** Heatmap of high entropy genes in all GBM samples in AOI from the combined tumor dataset. Highlighted genes are associated with cell types and discussed in the text. **d.** Expression of *APOE* in Visium® dataset in representative GBM samples from *Ruiz-Moreno* dataset[14]. **e.** Expression of *VEGFA* and Mes-like signature in representative GBM samples from Visium® dataset. Below, p-values of spatial correlation between VEGFA expression and mesenchymal signature.

### Supplementary figure 6

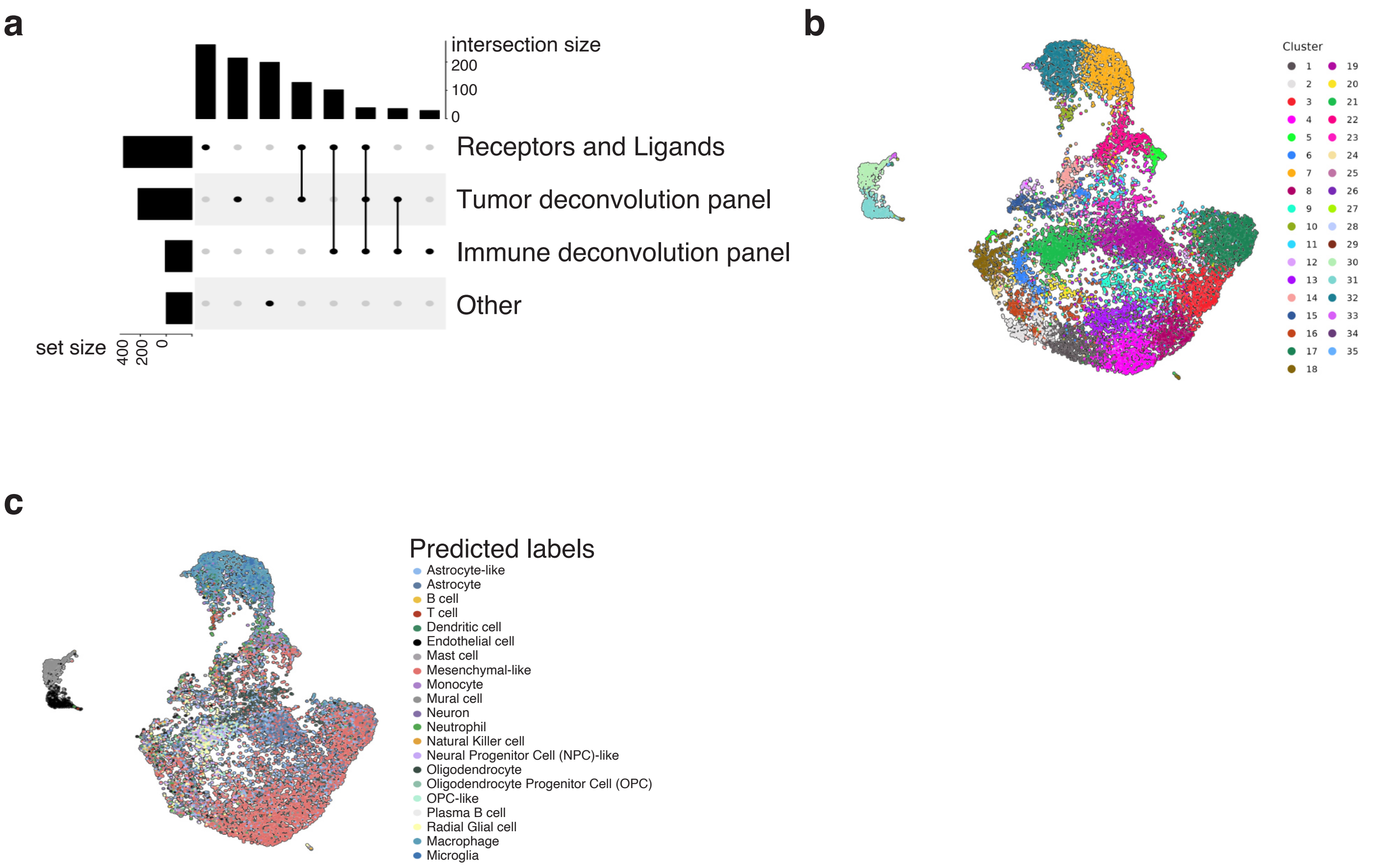

**Supplementary Fig. 6 |**

**a.** UpsetR plot of NanoString 1,000 onco-immunology panel. The number of probes represented in single signatures and the number of overlapping probes are visualized to generate the tumor and immune cell state signatures and for ligand/receptor analyses. **b.** UMAP plot of all 11,801 cells colored by cluster membership. **c.** UMAP colored by predicted cell types from harmonized GBM reference dataset[14].

### Supplementary figure 7

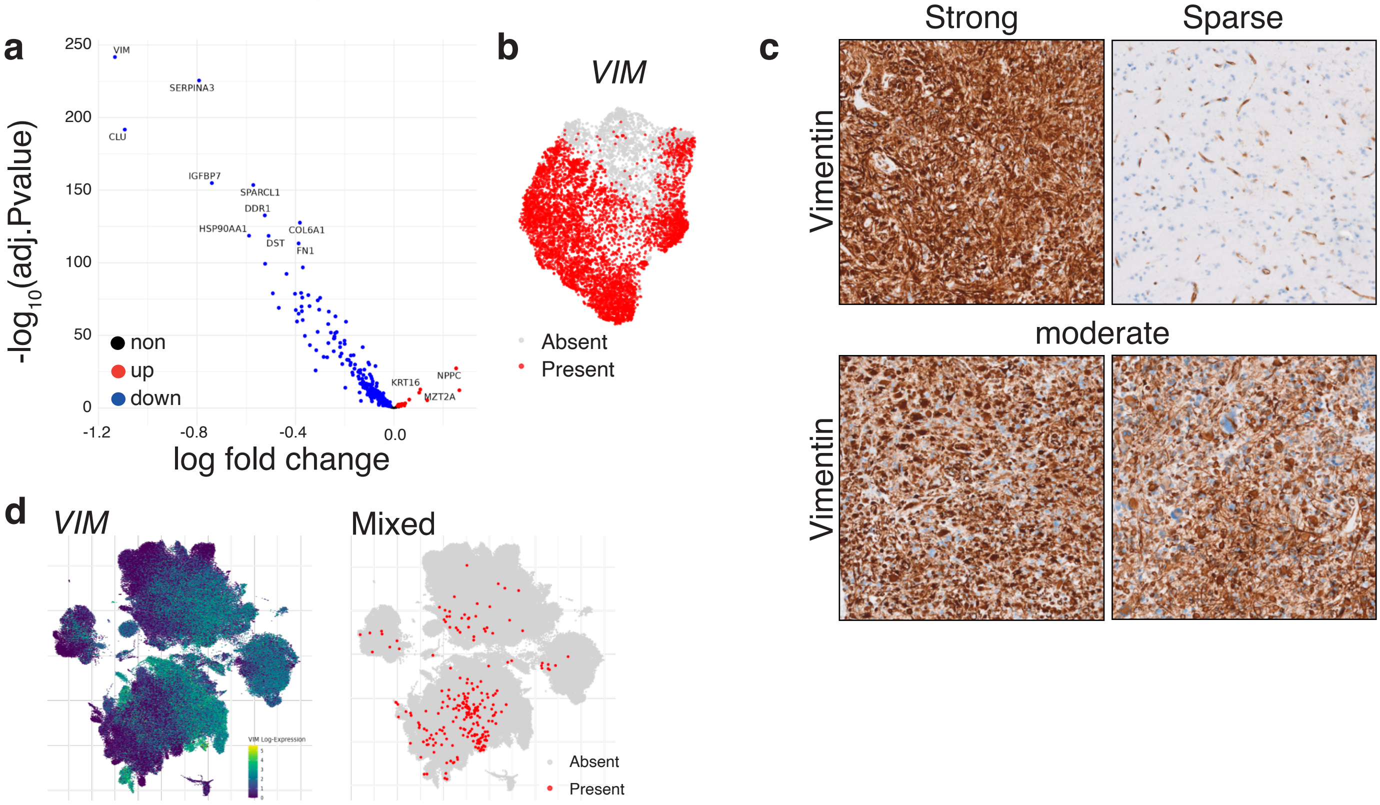


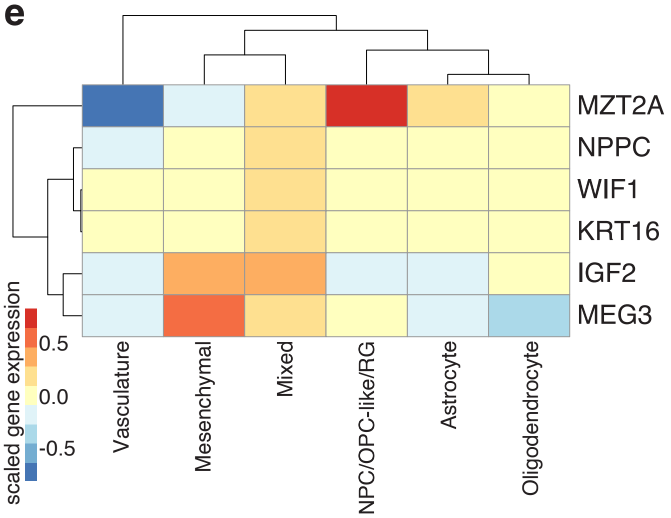


**f**

| Ontogeny | Term | P value |
|---|---|---|
| GO:0045236 | CXCR chemokine receptor binding | 0.00018 |
| GO:0061844 | antimicrobial humoral immune response mediated by antimicrobial peptide | 0.00024 |
| GO:0001664 | G protein-coupled receptor binding | 0.00029 |
| GO:0048018 | receptor ligand activity | 0.00059 |
| GO:0030546 | signaling receptor activator activity | 0.00074 |
| GO:0030545 | signaling receptor regulator activity | 0.00083 |
| GO:0005179 | hormone activity | 0.0010 |
| GO:0019730 | antimicrobial humoral response | 0.0018 |
| GO:0042379 | chemokine receptor binding | 0.0027 |
| GO:0070098 | chemokine-mediated signaling pathway | 0.0083 |

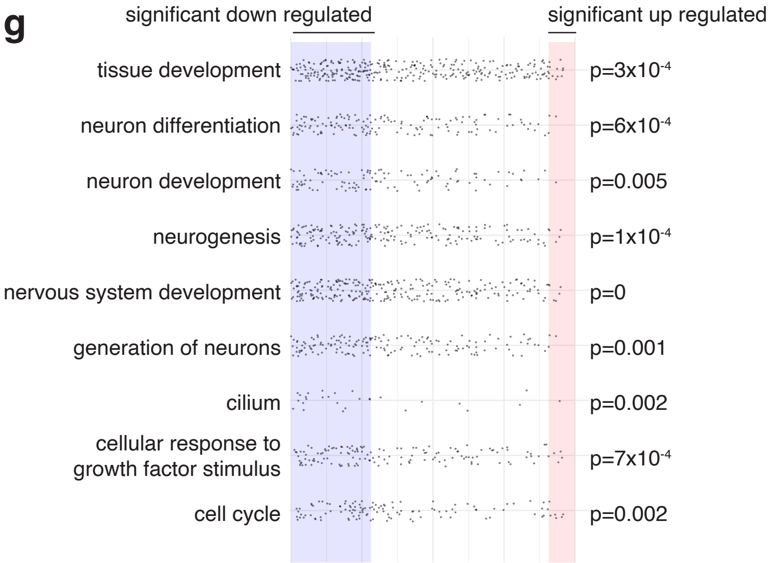

**Supplementary Fig. 7 |**

**a.** Volcano plot of differentially expressed genes in Mixed population compared to other tumor cell states; upregulated genes are in red and downregulated genes are in blue. **b.** UMAP plot of tumor cells indicating presence of *VIM*. **c.** Vimentin immunohistochemical staining across GBM-1 tumor. Representative examples of strong homogenous stain, sparse stain (in non-tumor region) and moderate staining of heterogeneous tumor regions. Scale, 100 µm. **d.** Expression of *VIM* (left) and location of nearest neighbors to the Mixed population in the *Ruiz-Mureno* dataset[14] (left). **e.** Heatmap of top up-regulated genes in Mixed population compared to other tumor populations. **f.** Table of top up-regulated gene ontology (GO) pathway terms in the Mixed population compared to other tumor populations. **g.** Dot plot displaying the rank of the respective genes of the top identified GO terms in the Mixed population compared to other tumor populations.

#### Supplementary figure 8

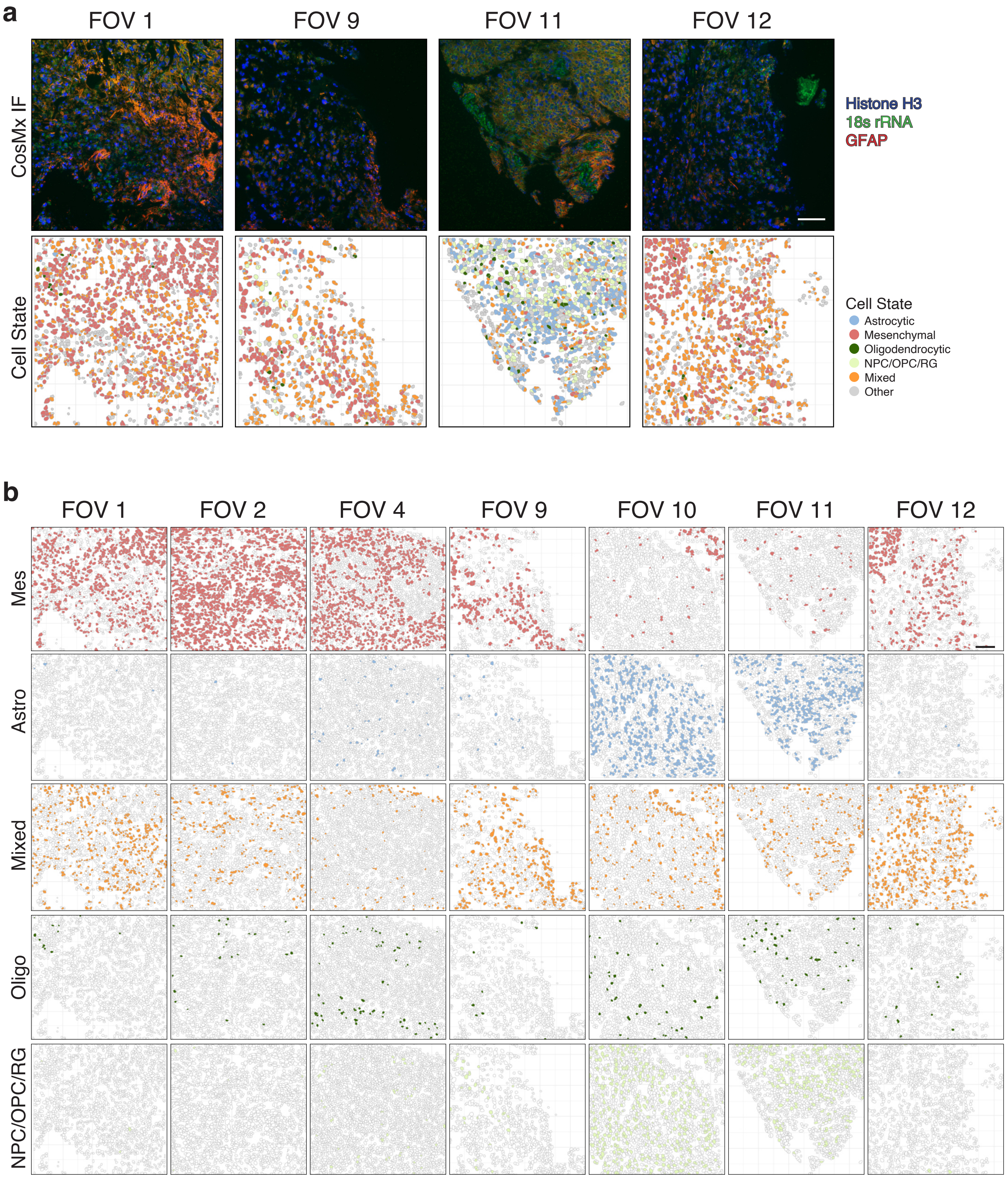

**Supplementary Fig. 8 |**

**a.** Panel of FOV 1, 9, 11, 12 CosMx® immunofluorescence protein expression (Histone H3, 18s rRNA, GFAP) and Voronoi plot of cell state assignment. Scale, 100 µm. **b.** Voronoi plots of cell state maps for each state and FOV. Scale, 100 µm.

### Supplementary figure 9

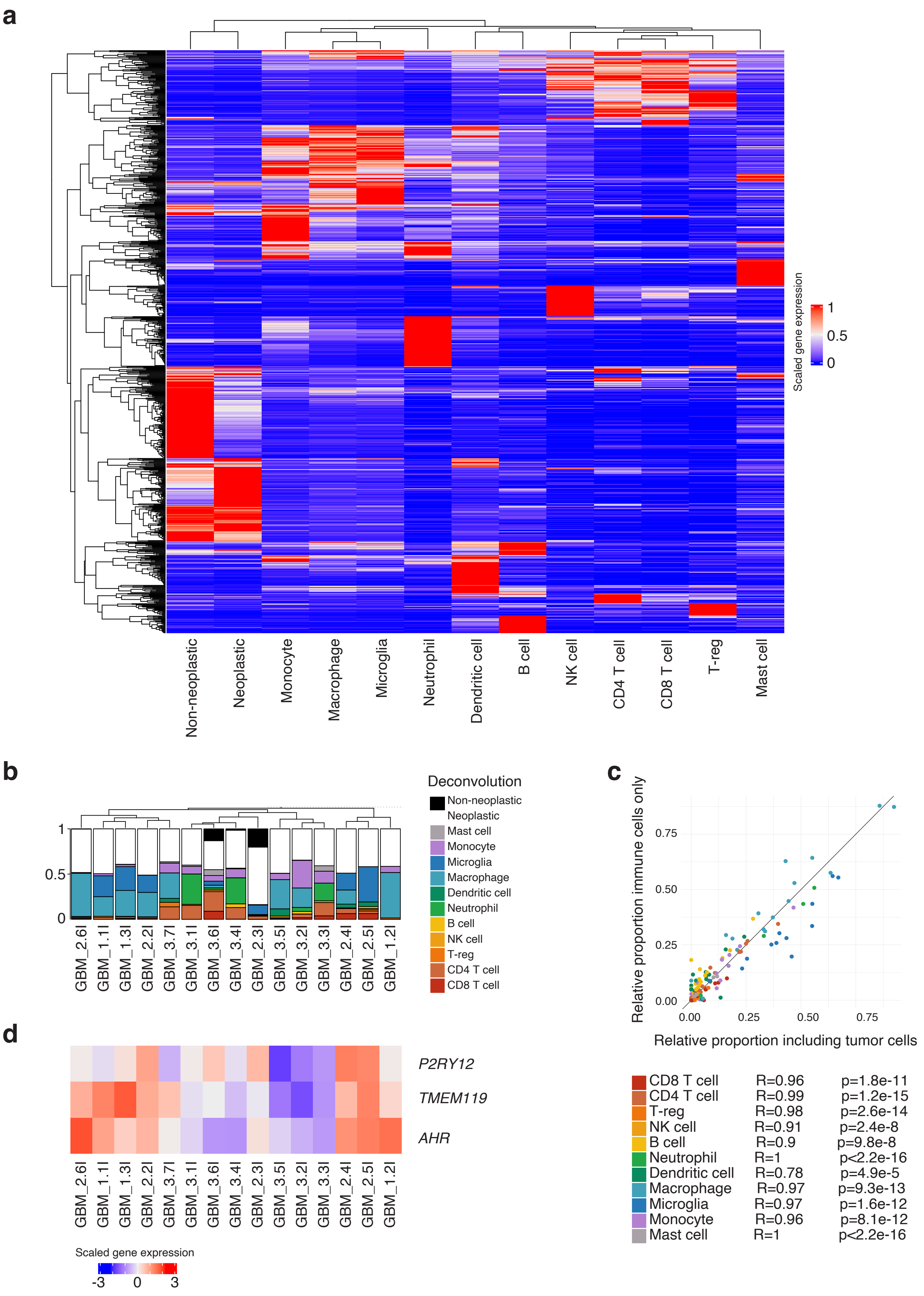

**Supplementary Fig. 9 |**

**a.** Heatmap of immune cell type deconvolution signatures. Scaled gene expression of genes from Ruiz-Moreno *et al.* dataset present in SafeTME reference and preprocessed gene list (n = 859), for each cell type profile. **b.** GBM Immune AOI deconvolution including non-immune neoplastic and non-neoplastic cell types. **c.** Immune proportions pre and post tumor cell deconvolution. Below, correlations of proportions for each cell type. **d.** Expression of key macrophage and microglia genes *P2RY12, TMEM119* and *AHR* in GBM immune AOIs.

### Supplementary figure 10

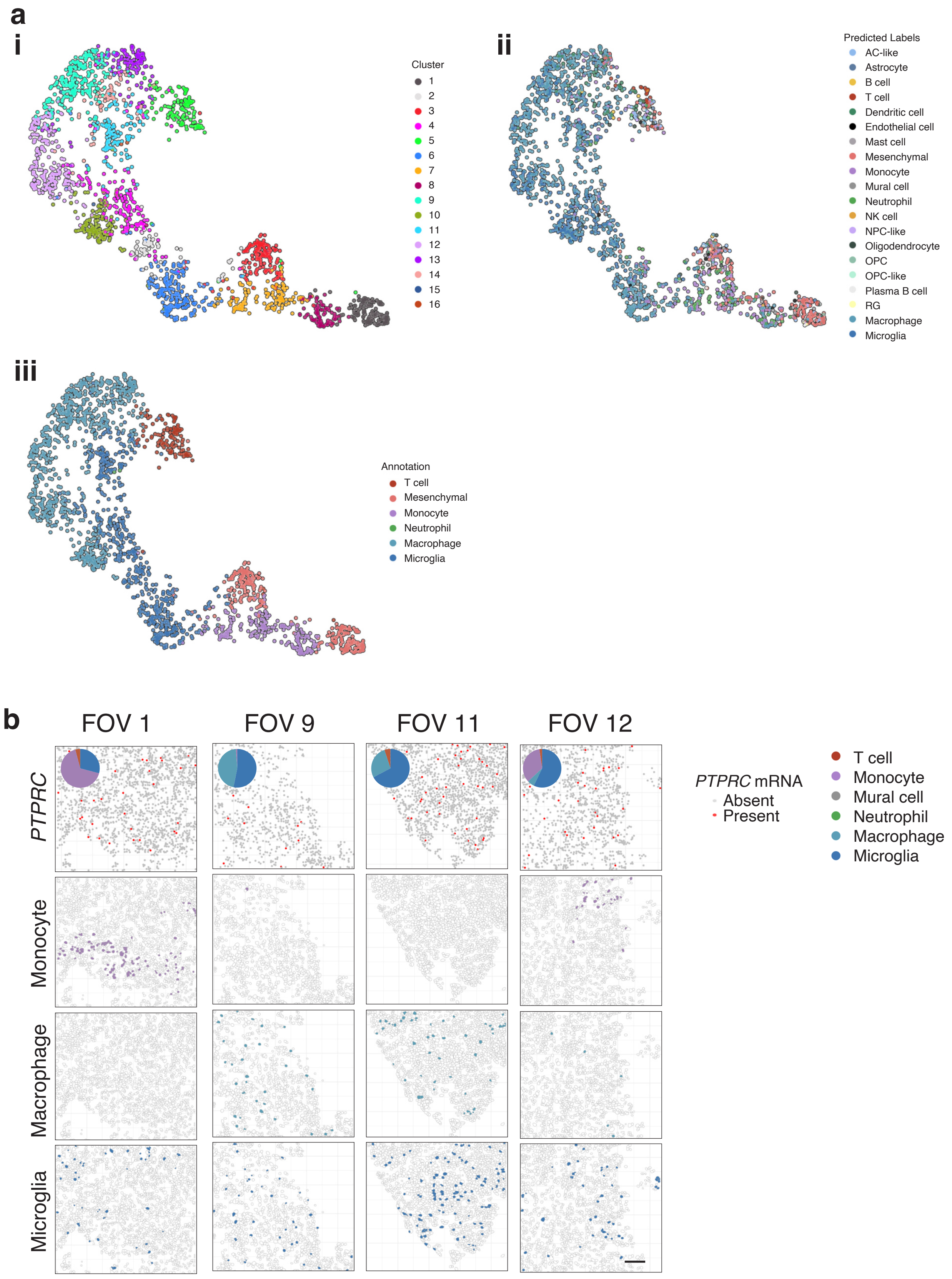

**Supplementary Fig. 10 |**

**a.** UMAP of 2,203 $CD45^+$ cells colored by (i) clusters (ii) predicted cell types and (iii) inferred cell states. **b.** Expression of *PTPRC* in spatial context with inset of immune infiltrate proportions; and Voronoi plots of monocyte, macrophage and microglia gene signatures in GBM-1 FOV1, 9, 11 and 12. Scale, 100 µm.

### Supplementary figure 11

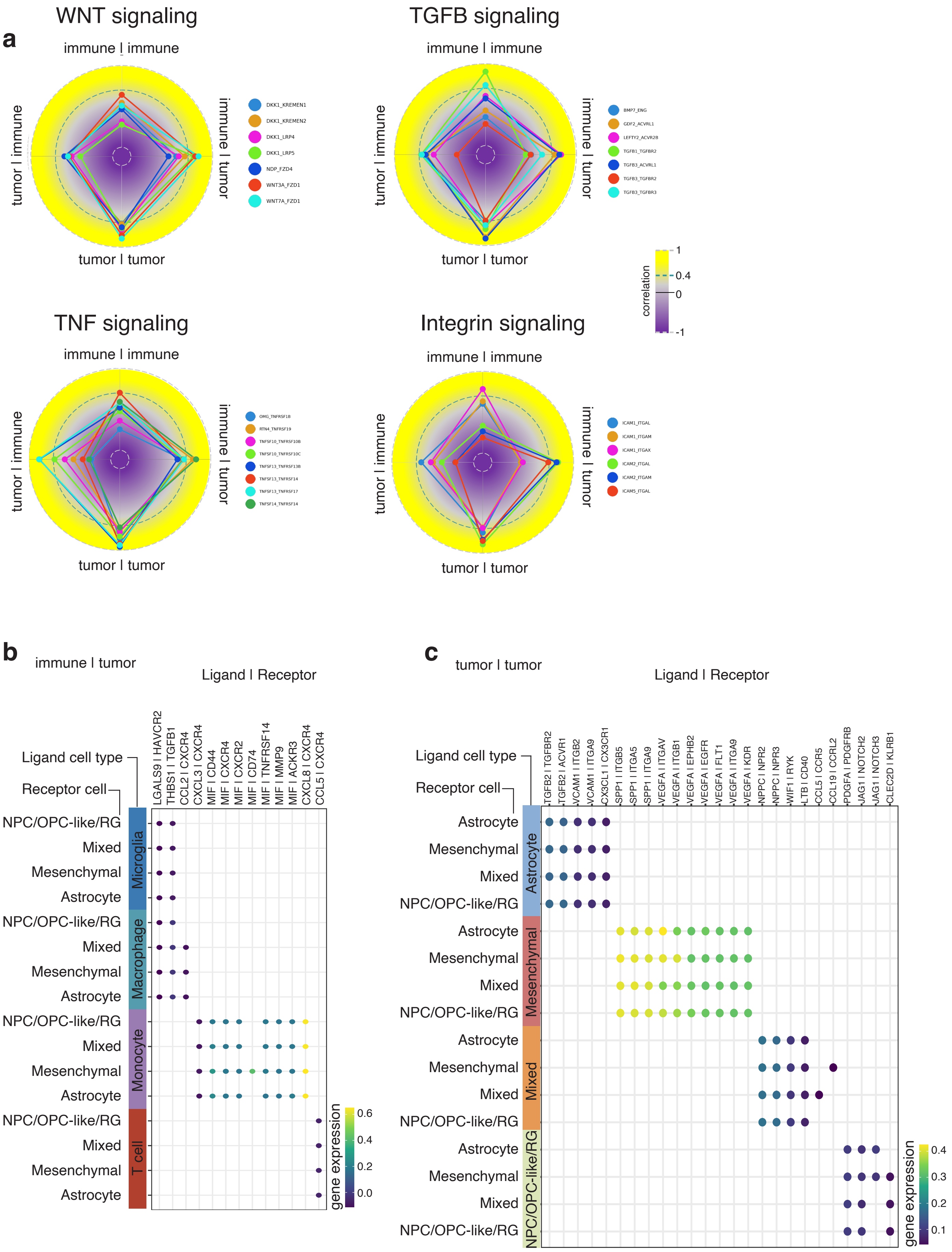

**Supplementary Fig. 11 |**

**a.** Correlation spider plots of ligand receptor pairs with pronounced tumor to tumor signaling. Top left, WNT signaling pathway; top right, TGFβ signaling pathway; bottom left, TNF signaling pathway; bottom right, ITG signaling pathway. **b.** Product of expression of significant top ligand/receptor interaction pairs identified from DSP between immune (ligand) and tumor (receptor) cells in GBM-1. **c.** Product of expression of significant ligand/receptor interaction pairs between tumor (ligand) and tumor (receptor) cells that show specificity with regards to tumor state (ligand) in GBM-1.

### Supplementary figure 12

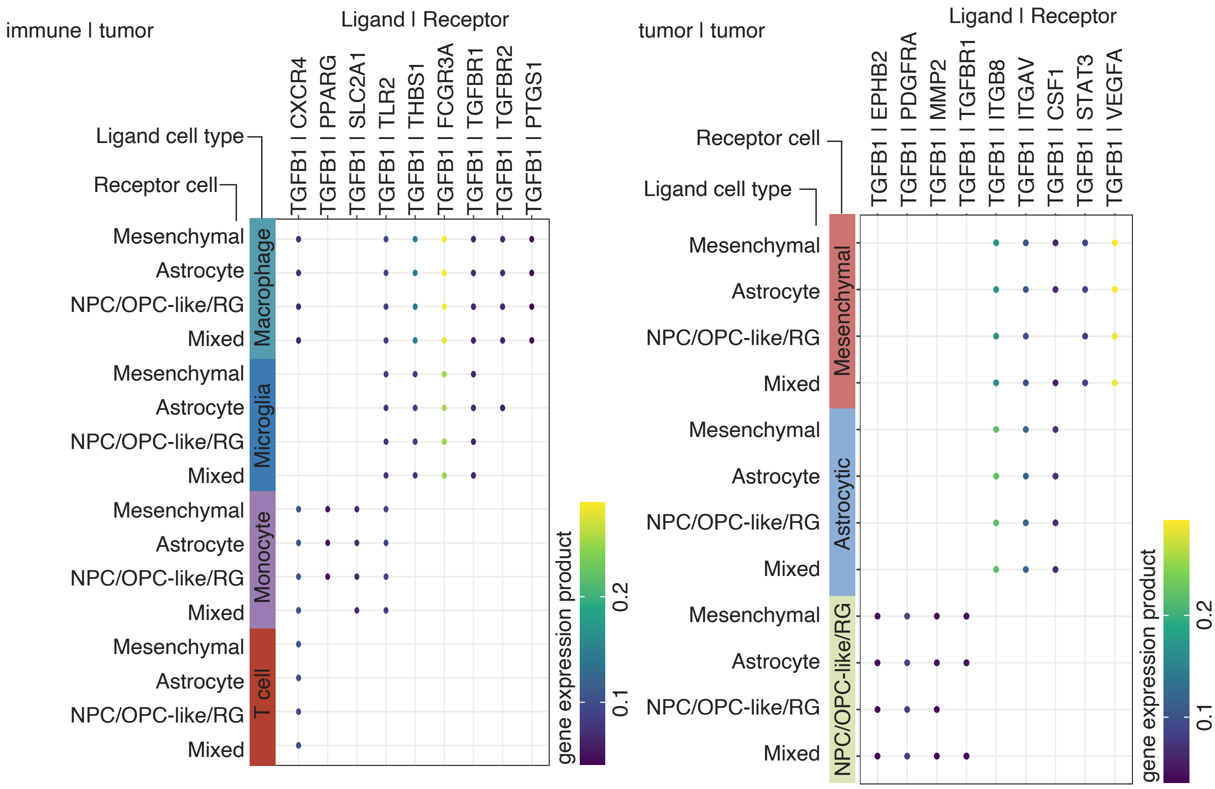

**Supplementary Fig. 12 |**

Product of expression of significant TGFB1 signaling ligand/receptor pairs in tumor-immune and tumor-tumor signaling.

#### Supplementary figure 13

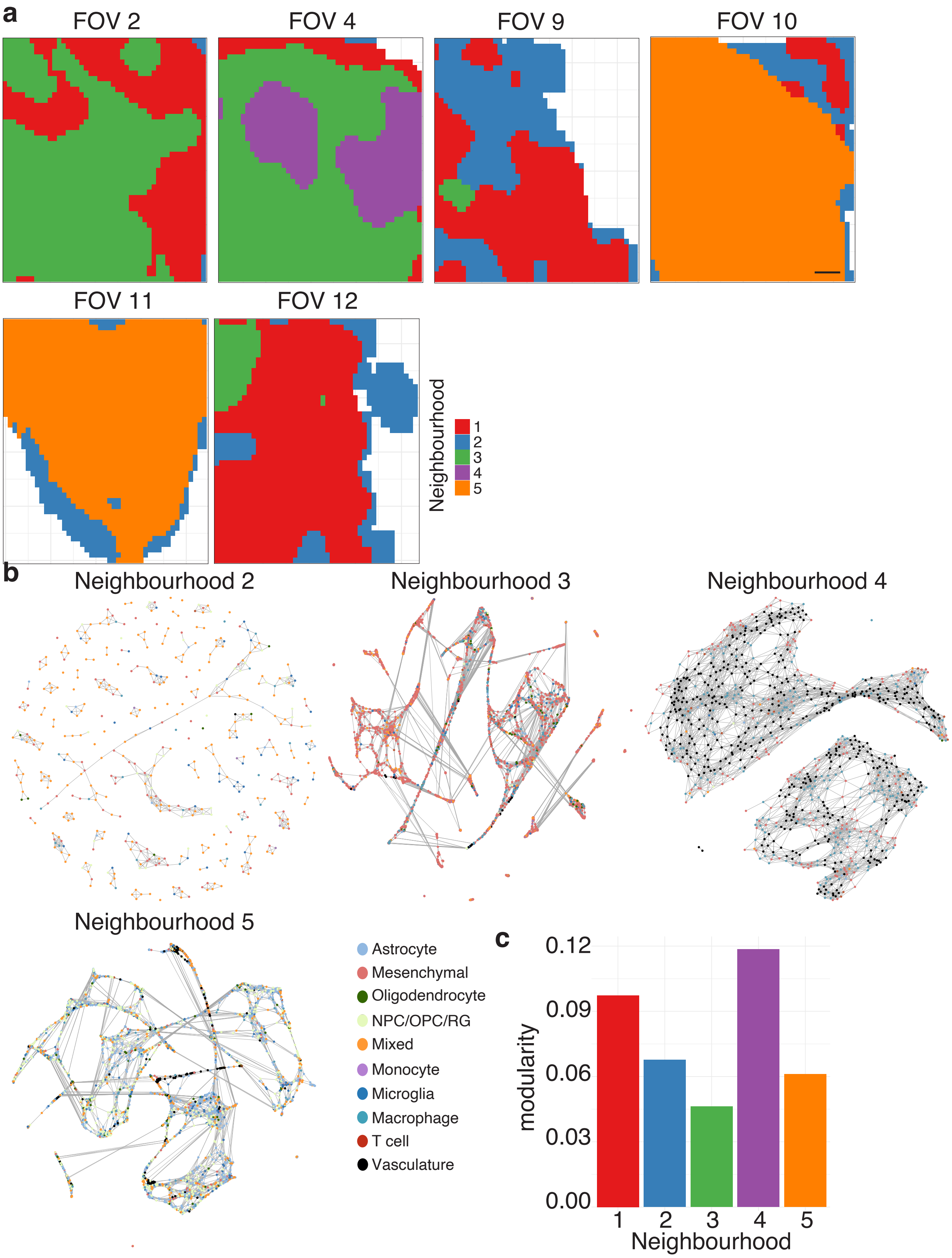

**Supplementary Fig. 13 |**

**a.** Plot indicating neighborhoods for each FOV in GBM-1 SMI analysis. Scale, 100 µm. **b.** Network displaying proximity of cell types within 30 µm radius in neighborhoods 2-5. **c.** Modularity score for each neighborhood measuring the number of connections between cells of the same state compared to those of other states.

#### Supplementary figure 14

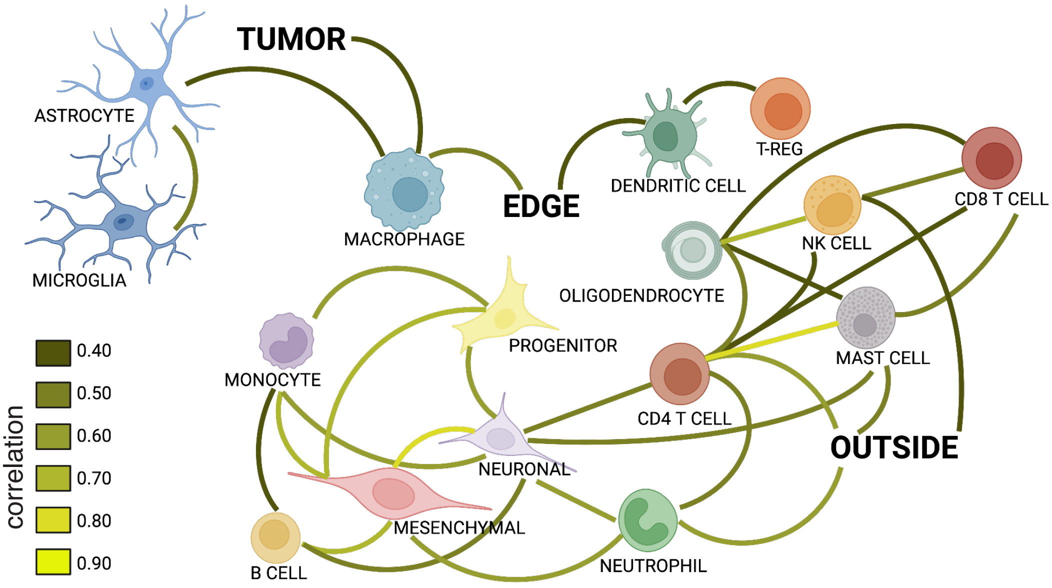

**Supplementary Fig. 14 |**

Schematic of cell co-occurrence in the key domains of the tumor, edge and outside from GeoMx® DSP data. Line color relative to correlation coefficient.

#### Supplementary Table 1: Astrocytoma samples AOI information

| Sample | ROI | Pathology Review | Primary Classification | Secondary Classification | Immune Classification | AOI | Segment | Characteristic | Status |
|---|---|---|---|---|---|---|---|---|---|
| A_1 | 001 | Cellular Tumor | Ki67 border | tumor | Low | GFAP+ | tumor | Ki67+ region | Included |
| | | | | | | CD45+ | immune | Ki67+ region | Excluded |
| | 002 | Cellular Tumor | tumor | | High | GFAP+Ki67+ | cycling tumor | tumor | Included |
| | | | | | | GFAP+ | non-cycling tumor | tumor | Included |
| | | | | | | CD45+ | immune | tumor | Included |
| | 003 | Cellular Tumor | tumor | | High | GFAP+Ki67+ | cycling tumor | tumor | Included |
| | | | | | | GFAP+ | non-cycling tumor | tumor | Included |
| | | | | | | CD45+ | immune | tumor | Excluded |
| | 004 | Cellular Tumor | tumor | | High | GFAP+ | tumor | necrotic | Included |
| | | | | | | CD45+ | immune | necrotic | Included |
| | 005 | Cellular Tumor | Ki67 border | | Low | GFAP+ | tumor | Ki67- region | Included |
| | | | | | | CD45+ | immune | Ki67- region | Excluded |
| A_2 | 001 | Cellular Tumor | Ki67 border | tumor | Low | GFAP+ | tumor | Ki67+ region | Included |
| | | | | | | CD45+ | immune | Ki67+ region | Excluded |
| | 002 | Cellular Tumor | Ki67 border | | Low | GFAP+ | tumor | Ki67- region | Included |
| | | | | | | CD45+ | immune | Ki67- region | Included |
| | 003 | Cellular Tumor | border | tumor | Low | GFAP+ | tumor | Ki67+ region | Included |
| | | | | | | CD45+ | immune | Ki67+ region | Excluded |
| | 004 | Infiltrating Tumor | border | | Low | GFAP+ | tumor | Ki67- region | Included |
| | 005 | Cellular Tumor | Ki67 border | tumor | Low | GFAP+ | tumor | Ki67+ region | Included |
| | | | | | | CD45+ | immune | Ki67+ region | Included |
| | 006 | Cellular Tumor | tumor | | High | GFAP+Ki67+ | cycling tumor | tumor | Included |
| | | | | | | GFAP+ | non-cycling tumor | tumor | Included |
| | | | | | | CD45+ | immune | tumor | Included |
| A_3 | 001 | Cellular Tumor | Ki67 border | | Low | GFAP+ | tumor | Ki67- region | Included |
| | 002 | Leading Edge | border | | High | GFAP+Ki67+ | cycling tumor | normal | Excluded |
| | | | | | | GFAP+ | non-cycling tumor | normal | Included |
| | 003 | Infiltrating Tumor | border | | High | GFAP+Ki67+ | cycling tumor | mix tumor | Included |
| | | | | | | GFAP+ | non-cycling tumor | mix tumor | Included |
| | 004 | Infiltrating Tumor | border | | High | GFAP+Ki67+ | cycling tumor | mix tumor | Included |
| | | | | | | GFAP+ | non-cycling tumor | mix tumor | Included |
| | 005 | Cellular Tumor | Ki67 border | tumor | Low | GFAP+ | tumor | Ki67+ region | Included |

#### Supplementary Table 2: Glioblastoma samples AOI information

| Sample | ROI | Pathology Review | Primary Classification | Secondary Classification | Immune Classification | AOI | Segment | Characteristic | Status |
|---|---|---|---|---|---|---|---|---|---|
| GBM_1 | 001 | Cellular Tumor and Microvascular Proliferation | tumor | | High | GFAP+Ki67+ | cycling tumor | tumor | Included |
| | | | | | | GFAP+ | non-cycling tumor | tumor | Included |
| | | | | | | CD45+ | immune | tumor | Included |
| | 002 | Cellular Tumor | tumor | | High | GFAP+Ki67+ | cycling tumor | tumor | Included |
| | | | | | | GFAP+ | non-cycling tumor | tumor | Included |
| | | | | | | CD45+ | immune | tumor | Included |
| | 003 | Cellular Tumor | tumor | | High | GFAP+Ki67+ | cycling tumor | tumor | Included |
| | | | | | | GFAP+ | non-cycling tumor | tumor | Included |
| | | | | | | CD45+ | immune | tumor | Included |
| | 004 | Cellular Tumor | border | tumor | Low | GFAP+ | tumor | tumor | Included |
| | 005 | Infiltrating Tumor | border | | Low | GFAP+ | tumor | mix tumor | Included |
| | 006 | Leading Edge | border | | Low | GFAP+ | tumor | normal | Included |
| GBM_2 | 002 | Cellular Tumor | border | | Low | GFAP+Ki67+ | cycling tumor | tumor | Included |
| | | | | | | GFAP+ | non-cycling tumor | tumor | Included |
| | | | | | | CD45+ | tumor | tumor | Included |
| | 003 | Infiltrating Tumor | border | | Low | GFAP+Ki67+ | cycling tumor | normal | Excluded |
| | | | | | | GFAP+ | non-cycling tumor | normal | Included |
| | | | | | | CD45+ | tumor | normal | Included |
| | 004 | Cellular Tumor | Ki67 border | tumor | High | GFAP+ | tumor | Ki67+ region | Included |
| | | | | | | CD45+ | immune | Ki67+ region | Included |
| | 005 | Cellular Tumor | Ki67 border | | High | GFAP+ | tumor | Ki67- region | Included |
| | | | | | | CD45+ | immune | Ki67- region | Included |
| | 006 | Cellular Tumor | border | | Low | GFAP+Ki67+ | cycling tumor | tumor | Included |
| | | | | | | GFAP+ | non-cycling tumor | tumor | Included |
| | | | | | | CD45+ | tumor | tumor | Included |
| GBM_3 | 001 | Pseudo-palisading cells around Necrosis | Ki67 border | | Low | GFAP+ | tumor | Ki67- region | Included |
| | | | | | | CD45+ | immune | Ki67- region | Included |
| | 002 | Cellular Tumor | Ki67 border | tumor | Low | GFAP+ | tumor | Ki67+ region | Included |
| | | | | | | CD45+ | immune | Ki67+ region | Included |
| | 003 | Perinecrotic Zone | Ki67 border | | Low | GFAP+ | tumor | Ki67- region | Included |
| | | | | | | CD45+ | immune | Ki67- region | Included |
| | 004 | Pseudo-palisading cells around Necrosis | Ki67 border | | High | GFAP+ | tumor | Ki67- region | Included |
| | | | | | | CD45+ | immune | Ki67- region | Included |
| | 005 | Pseudo-palisading cells around Necrosis | Ki67 border | tumor | High | GFAP+ | tumor | Ki67+ region | Included |
| | | | | | | CD45+ | immune | Ki67+ region | Included |
| | 006 | Necrosis | border | | High | GFAP+ | tumor | normal | Included |
| | | | | | | CD45+ | immune | normal | Included |
| | 007 | Cellular Tumor | Ki67 border | tumor | Low | GFAP+ | tumor | Ki67+ region | Included |
| | | | | | | CD45+ | immune | Ki67+ region | Included |
